# Interactions between epistasis and pleiotropy in generic models of complex traits

**DOI:** 10.64898/2026.09.29.755220

**Authors:** Caelan Brooks, Misha Gupta, Gautam Reddy, Michael M. Desai

## Abstract

Evolution in fluctuating environments depends critically on epistatic interactions between mutations and the pleiotropic effects these mutations have on fitness across multiple conditions. However, while these effects of epistasis and pleiotropy have been extensively studied individually, the interactions between them are less well understood. In this study, we analyze how genetic background shapes the effects of mutations across multiple environmental conditions in generic models of complex traits. We show that when epistasis is widespread and random, systematic patterns of cross-environment pleiotropy can emerge naturally. Specifically we outline two regimes, *widespread random epistasis* (WRE) and *structured pleiotropy* (SP), in which patterns of global epistasis in one environment can be predicted from those in other conditions. We outline these predictions in several classes of generic null models, and test these predictions in laboratory budding yeast by measuring the fitness effects of a large set of mutations across many genetic backgrounds in several growth environments. We find that only a subset of mutations fall within the WRE and SP regimes, but that in these cases two of our models provide accurate fits to the data. Finally, we use this framework to show how modular, pleiotropic architectures shape long-term evolutionary steady states in fluctuating environments, revealing how tradeoffs between environments can develop from the geometry of shared latent interaction structure.

## Introduction

Epistatic interactions between mutations play a critical role in evolutionary adaptation. If epistasis is rare, then the effects of new mutations are largely independent of the genetic background in which they occur, and the fitness landscape is relatively smooth. In this case, information about the effects of single substitutions on a few genetic backgrounds is sufficient to characterize the landscape, and predicting evolutionary dynamics is relatively straightforward. On the other hand, if epistasis is widespread, then the effects of a given mutation can change dramatically across genetic backgrounds, the fitness landscape becomes rugged, and evolutionary dynamics can become much more complex.

Numerous empirical studies have shown that epistasis is indeed widespread among mutations that accumulate during microbial adaptation, both in laboratory evolution experiments [Bakerlee et al., 2022, Lenski et al., 2015, Perfeito et al., 2014, Couce and Tenaillon, 2015] as well as in natural populations [Moulana et al., 2022, Witte et al., 2023, DelaFuente et al., 2024, Bourg et al., 2024]. In principle, this epistasis could be arbitrarily complex, reflecting the idiosyncratic biological details of each specific organism and environmental context. However, recent work has shown that in many microbial systems, a general pattern of “global” diminishing-returns and increasing-costs epistasis emerges: beneficial mutations become systematically less beneficial and deleterious mutations become systematically more deleterious as the fitness of the genetic background in which they occur increases [Kryazhimskiy et al., 2014, Good and Desai, 2015, Diaz-Colunga et al., 2023, González-González et al., 2024]. Rather than making evolution less predictable, this leads to an overall pattern of declining adaptability (i.e. the rate of fitness increase slows down as populations adapt), which leads to reproducible patterns of convergent evolution (at the level of fitness changes over time) across replicate populations [Wiser et al., 2013, Good et al., 2017, Johnson et al., 2021].

Global patterns of diminishing returns and increasing costs epistasis could in principle reflect physiological constraints or specific features of the structure of genetic networks. However, recent theoretical work by us [Reddy and Desai, 2021] and by others [Lyons et al., 2020] has shown that these patterns emerge as a generic consequence of large numbers of random epistatic interactions. In other words, global epistasis may not reflect any particular biological feature or structure, but instead can emerge naturally from many random and unstructured interactions. This provides a natural null model for the role of epistasis in microbial adaptation, which makes testable predictions that we have explored in previous work [Reddy and Desai, 2021].

Prior analysis of the implications of these generic models of complex traits has focused on evolution in a single environment. When environmental conditions can change across time or space, the pleiotropic effects of mutations on fitness in multiple environments also become critical. Extensive prior work [Martin and Lenormand, 2015, Kinsler et al., 2020, Geiler-Samerotte et al., 2020, Wang et al., 2024] has characterized pleiotropy among individual mutations (e.g. finding patterns of antagonistic pleiotropy that reflect fundamental physiological tradeoffs, or systematic positive correlations in the effects of mutations on similar phenotypes). However, the interactions between epistasis and pleiotropy (i.e. how the genetic background influences the effects of mutations across multiple environments) are less well understood. These interactions determine how evolution in one environment affects the fitness effects of potential new mutations in other conditions, and hence are critical for understanding evolution in changing environments. One recent study has argued that global epistasis is nearly invariant across different phenotypes (specifically, growth rates in different environmental conditions), and is modulated only by a single environment-specific “pivot” growth rate [Ardell et al., 2024]. This provides clear expectations for how evolution in one condition affects potential further adaptation in other environments. However, the generality and biological basis of this observation is not known.

Here, we analyze the interactions between epistasis and pleiotropy in generic models of complex traits, generalizing the single-trait random landscapes model considered by Reddy and Desai [2021]. We explore several simple null expectations for the structure of these random models. We find that systematic patterns of pleiotropy emerge naturally, provided that epistatic and pleiotropic effects are random and sufficiently widespread. To test these predictions, we created a large library of specific gene disruption mutations across a panel of budding yeast genetic backgrounds, and measured the effects of these disruptions on a set of five growth rate phenotypes. We find that the assumptions of our generic null models only hold for a subset of these mutations, but that for this subset our models provide an excellent fit to the experimental data. This suggests that systematic patterns in the interactions between epistasis and pleiotropy in this system need not reflect idiosyncratic details of the biology, but instead can (in at least a subset of cases) simply emerge as the consequence of many random interactions shared across environments. Finally, we explore the consequences of these random models of complex traits for evolutionary dynamics and long-term steady states in fluctuating environmental conditions.

## Analysis and Results

### Model

Results in Reddy and Desai [2021], establish that patterns of global epistasis can result from many microscopic interactions in a single environment. The magnitude of the relationship between a mutation’s fitness effect and the genotype’s fitness before the mutation is made reflects how interactive a locus is, and a large number of interactions predicts a specific link between the slope and variance of this relationship. In this study, we ask whether many interactions across environments give rise to global structure, and whether that structure becomes predictive when the interactions are shared across environments.

We consider an experimental setting in which we measure the fitness effect of a mutation at each of *F* “focal” loci in some set of *E* different environments (or more generally we measure some set of *E* different phenotypes). We do so across a large number of genetic backgrounds, which vary across a set of *L* biallelic “background” loci. We write the genotype at the background loci as 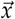, where *x*_*i*_ ∈ [ −1, 1]. We denote the genotypes at the focal loci as *xα* (note we reserve latin subscripts for the *L* background loci and greek subscripts for the *F* focal loci). We define the wild-type state of each focal locus to be *x*_*α*_ = −1, and the mutated state is *x*_*α*_ = +1.

We begin by analyzing the case of a single focal locus (i.e. *F* = 1), which illustrates the key ideas in the simplest possible context. In the Supplementary Text, we describe how to generalize these results to multiple focal loci, which is conceptually similar but introduces some technical complications. In this *F* = 1 case, we write a general expression of our *E* genotype-phenotype landscapes,

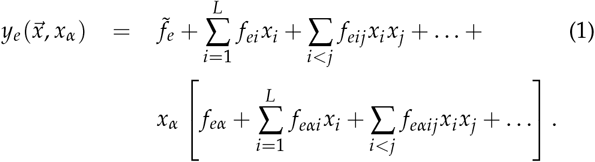

Here *y*_*e*_ is the fitness in environment *e* (or more generally phenotype *e* from among *E* total measured phenotypes), 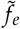 is the mean fitness in that environment (averaged across all possible genotypes at the background and focal loci, assuming −1 and +1 have equal probabilities), the *f*_*ei*_ and *f*_*eα*_ terms are the background-averaged linear effect of each locus in that environment, the *f*_*eij*_ and *f*_*eαi*_ terms represent background-averaged pairwise epistatic interactions, *f*_*eijk*_ and *f*_*eαij*_ terms represent third-order epistasis, and so on. We emphasize that this formulation of the genotype-phenotype map is entirely general, and we have not yet introduced any assumptions about the structure of the landscape. In other words, any genotype-phenotype map involving *L* + 1 biallelic loci can be written in this form.

We will find it convenient to introduce the definitions

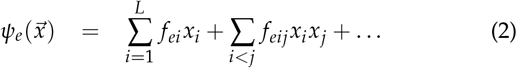

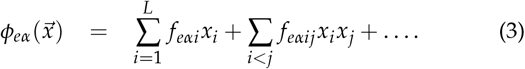

With these definitions, we have

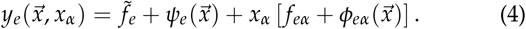

We note that by construction we will have ⟨*ψ*_*e*_⟩ = ⟨*ϕ*_*eα*_⟩ = 0, where ⟨·⟩ denotes an average over genotypes at the background loci. It is useful to also define 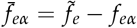.

In our typical experimental setting, we will have measurements of 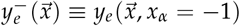 and 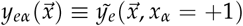, for each of a large number of genotypes at the background loci 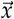. We note that for *n* such background genotypes, we have 2*nE* total measurements plus 2*E* constraints, from which we can compute the (2*n* + 2)*E* quantities 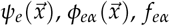, and 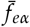.

### Cross environment global epistasis

The fitness effect of the mutation at the focal locus in environment *e* and genetic background 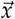 is defined as 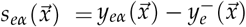. We begin by writing out how this expected fitness effect *s*_*eα*_ depends on the background fitness in another environments, 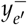:

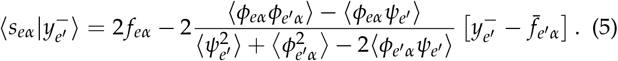

This result is a general feature of linear regression; we have not yet made any approximations. We note that the slope of the dependence of the fitness effect in environment *e* on the background fitness in environment *e*^′^ depends on relationships between the interaction terms in the two environments, including both those that involve the focal locus and those that only involve background loci. Specifically, we can think of 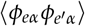 (which involves sums of terms of the form 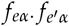) as summarizing the relationships (across environments *e* and *e*^′^) between interaction terms involving the focal locus. In contrast, the 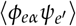 covariances involve sums of terms of the form 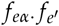, which involve mixed orders of epistasis (an order *q* term involving the background loci times an order *q* + 1 term involving those background loci and the focal locus). In the sections below, we consider how we expect these quantities to be related in various types of genotype-phenotype landscapes.

### The widespread random epistasis (WRE) and structured pleiotropy (SP) limits

In principle, the quantities 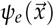 and 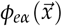 can have arbitrarily complicated dependence on the background genotype 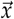. However, we focus in this work on generic models of complex traits in which epistasis is both widespread and random. By widespread, we mean that there are a large number of nonzero epistatic *f* terms (i.e. at second order or above). By random, we mean that there are no structured relationships among terms that reflect interactions between different sets of loci (i.e. different *f* terms are independent), though we do allow for relationships between the same term in different environments (these will reflect patterns of pleiotropy).

To be more precise, we focus on the *widespread random epistasis* (WRE) limit from Reddy and Desai [2021], in which we can make two key simplifications. First, we assume that we can treat the 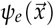 and 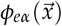 as jointly Gaussian, which will be true provided that these terms involve sums over a sufficiently large number of independent random variables that the central limit theorem applies. Second, we assume that expectations of the form ⟨*ψϕ⟩* are small compared to those of the form ⟨*ψ*^2^⟩ and ⟨*ϕ*^2^⟩. We might expect this to be true because the ⟨*ψ*_*e*_*ϕ*_*eα*_⟩ terms involve sums over products of different *f* terms (which average to 0) while the 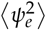 and 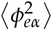 terms involve sums over squares of *f* terms (which are always positive). Therefore, the former will typically have a magnitude that is smaller by the square root of the number of terms in each sum. However, we note that the terms in the sums involved in the former sums are on average at lower epistatic order, so roughly speaking this condition will only hold provided that the sparsity or magnitude of epistatic terms falls off more slowly than as the square root of the epistatic order (though cutoffs such that interaction terms vanish above some specified order are not necessarily inconsistent with the WRE limit). More precise details describing these approximations and the specific conditions under which they apply can be found in Reddy and Desai [2021].

The WRE limit focuses on the relationship among expectations involving *ψ* and *ϕ* in a single environment. For our analysis of pleiotropy, we must also consider cross-environment expectations. Towards this purpose, we also define the *structured pleiotropy* (SP) limit. This SP limit is a cross-environment generalization of the WRE, in which we assume that 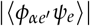 is small compared to 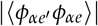. This requires that sums of terms of the form 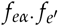 are small compared to sums of the form 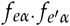. This is similar to the requirement for the WRE limit, but now involving products of interaction terms in the two different environments. We expect correlations between environments which involve products of *f* values of the same order to be larger than those involving products of *f* values of different order, and therefore this limit to hold for the same reason as the WRE limit. However, the conditions of validity are more stringent, because the overall magnitude of the sums of the latter terms is now scaled by the correlation between interaction terms across environments (which by definition is always less than or equal to 1).

The widespread epistasis limit represents a natural generic model of a complex trait that is determined by numerous random epistatic interactions. The structured pleiotropy limit extends this to settings where these random interactions have sufficiently strong correlations across environments (i.e. when there is some overall pattern of pleiotropy). We emphasize that neither the WRE or SP limits are guaranteed to hold for any particular focal mutation or set of background genotypes and environments. However, we can use experimental data to directly assess whether one or both limits holds in each specific case, as described below.

### Global epistasis and pleiotropy in the WRE and SP limit

In the WRE and SP limits, Eq. (5) becomes

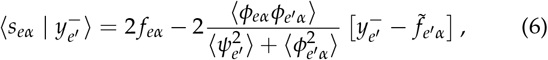

and the variance around this mean value is given by

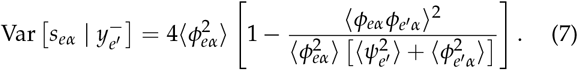

It will be helpful to consider how the expectations over *ψ*_*e*_ and *ϕ*_*eα*_ depend on the underlying landscape. We define

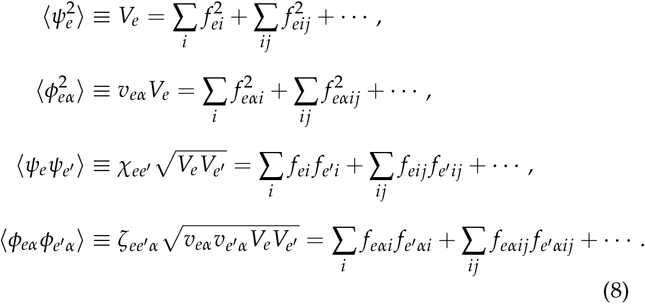

Here we have introduced the quantities 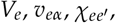 and 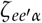 for future convenience (note that at this stage there is no approximation involved; we have defined *E*^2^ + *E* quantities to describe the *E*^2^ + *E* expectations). Roughly speaking, we can think of *V*_*e*_ as describing the total variance in fitness in environment *E* due to the background loci, *v*_*eα*_ as the fraction of this background variance that is added by terms involving the focal locus, 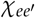 as a measure of the correlation in *f* terms among the background loci between environments *e* and *e*^′^, and 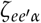 as a measure of this correlation among the *f* terms involving the focal locus. Note that by definition *V*_*e*_ and *v*_*eα*_ are always positive, while 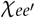 and 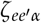 can be either positive or negative.

We note that in the typical experimental setting we envision, we can compute all the expectations described above directly from the data, and hence estimate 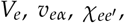, and 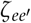. We also note that we can compute expectations of the form ⟨*ψ*_*e*_*ϕ*_*eα*_ ⟩ and 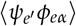, and assess whether thes e ex pe cta tions are indeed small compared to those of the form ⟨*ψ*^2^⟩,⟨*ϕ*^2^⟩ and 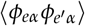, and therefore whether the WRE and SP limits are appropriate for a given focal locus and set of environments.

Using these definitions, we can see that when *e* = *e*^′^ we have

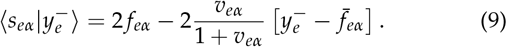

This is identical to the result in Reddy and Desai [2021], and represents a general pattern of global diminishing returns (or increasing costs) epistasis: the mutation at the focal locus becomes less beneficial (or more deleterious) in a given environment as the fitness of the genetic background it is made in increases. However, the dependence of the fitness effect of the focal mutation in one environment on the background fitness in the other environment is more complex; we have

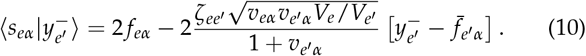

This dependence can be either positive or negative, since 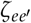 can have either sign. This means that as a population adapts in one environment, mutational effects in other environments can in principle either increase or decrease. However, we consider below two natural null models where this relationship can become predictable (fully random pleiotropy and the connectedness model), as well as one prediction related to the assumption of global epistatic structure (“criss-cross” environment predictions).

### Empirical tests of our generic null models

To test the conditions of the WRE and SP limits as well as the predictions of our generic models of complex traits outlined below with empirical data, we implemented the experimental system we envisioned above, using a set of 91 focal mutations in each of 87 genetic backgrounds. We then measured background fitnesses and the fitness effect of each focal mutation in each background in each of 5 different environmental conditions. The resulting data was then used to test the predictions of our generic null models above. Broadly speaking, we find that the WRE and SP approximations are only appropriate for a modest fraction of the focal mutations and environments we consider. However, in the cases where these approximations are valid, a modular model model provides a good fit to the experimental data (while a fully random pleiotropy model generally does not). We also find that predictions using cross environment global epistasis provide a good fit to the data even when the WRE and SP approximation do not apply.

### Measuring empirical patterns of epistasis and pleiotropy

To implement our empirical tests, we made use of a laboratory *Saccharomyces cerevisiae* strain collection constructed by Johnson et al. [2019], in which a set of 96 mutations (*F* = 91 focal mutations plus 5 neutral controls) was introduced into each of *n* = 176 genetic backgrounds (these backgrounds were haploid F1 segregants from a cross between a laboratory strain BY and a wine strain RM, which differ at *L* ≈ 42, 000 background loci). Each such strain was constructed in several replicates, and each replicate of each strain was labeled with a unique DNA barcode.

For this work, we focused on a subset of *n* = 87 of these genetic backgrounds, and measured the fitness effect of each focal mutation and neutral control in *E* = 5 different growth environments using bulk barcode-based fitness assays. We then used flow cytometry-based assays to independently measure the fitness of each genetic background in each environment. Details of media conditions for each environment, as well as our protocols for barcode and flow-cytometry based fitness assays and our methods for fitness inference are described in Materials and Methods.

Using this approach, we measured 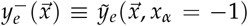 and 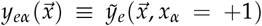, for each of our 91 focal loci *α* in each of our 5 environments *e* and 87 genetic backgrounds 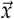. This represents a total of 39, 585 fitness measurements, although 12, 100 measurements dropped out (see Supplemental Information). From these measurements, we compute 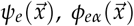, and *f*_*eα*_ for each focal mutation and environment. Note that in each of these computations, we use the correction for multiple focal loci as described in the Supplementary Text to compute the relevant quantities for each focal mutation (so for example 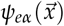 is different for each focal mutation).

### Testing the WRE and SP approximations

We first used this empirical data to compute ⟨*ϕ*_*eα*_*ψ*_*e*_ ⟩ and 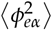 for each focal mutation and environment. We can then compare these quantities to assess whether the WRE approximation holds. Similarly, for each focal mutation and pair of environments we compute 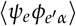 and 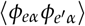, and compare these quantities to assess whether the SP approximation holds. We find that slightly more than half (58% on average over all environments) of the focal mutations in each environment fall in the WRE limit (defined as having 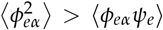; see Fig. S6). This is consistent with the result from Reddy and Desai [2021] using data from Johnson et al. [2019], which also found that the WRE applied for about half of the focal mutations they analyzed. As expected, somewhat fewer mutations (about 23% when averaging over all environment combinations) also fall within the more stringent SP limit (defined as those for which 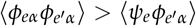; see Fig. S7).

The fact that only a modest fraction of focal mutations fall within the WRE and SP limits demonstrates that the assumptions of our generic null models are often violated. For example, it may be natural to expect that the magnitude of higher-order interaction terms often decays more quickly than the square root of the order (or that the sparsity of interactions increases more quickly than this). In this case, we do not expect the WRE or SP limits to generically hold. There may also be non-random structure in the epistatic terms for individual mutations, and in the case of the SP limit the correlation in interaction terms across environments may be too weak for sufficient pleiotropic structure to be present. These violations may be interesting in their own right, and point to focal mutations with nonrandom structure that merits further analysis. However, we do not expect the predictions from our analysis of generic models to hold in these cases (though the “criss-cross” predictions described below may be more generally valid). Nevertheless, we believe that these null models represent a useful generic baseline, and in comparisons with data below we highlight the focal mutations and environments which satisfy the WRE and SP assumptions, for which we expect our analysis to apply.

### Fully random pleiotropy model

#### Model

The first model we consider is a natural null model in which there is nothing “special” or idiosyncratic about the focal locus. In this case, we expect that the statistics of the relationship between *f* terms in the two different environments are the same regardless of whether the focal locus is involved, and hence we expect 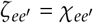. We refer to this as a model of “fully random pleiotropy.” In this fully random case, we can fit the 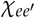 terms by considering the correlations in fitness across environments among the background genotypes and ask whether this correlation is representative of the relationship between focal mutations across those same environments.

#### Experimental test

To test whether the cross-environment correlations in the interaction terms involving the focal locus are similar to those among the background loci, we compute 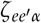 and 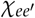 from the data (using the formulas defined in Eq. (8)) for each focal mutation and environment-pair, and determine whether they are equal.

In Fig. 1 we show the results of this test for all focal mutations in six of the ten possible environment-pairs (results for the other four environments are shown in Fig. S8). We see that while this model is appropriate for a few focal mutations (those that fall on the dotted one-to-one line), it fails in the majority of cases. We also note that, with a few exceptions, background fitness values as well as focal mutational effects tend to be positively correlated between our environments 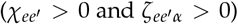. There is also more variation in how focal mutations are correlated compared to backgrounds. Together, these results indicate that pleiotropic structure is typically more complex than captured by the fully random pleiotropy model (even for focal mutations in the WRE and SP limits), and that considering how correlations in interaction terms differ across loci is important.

**Figure 1:**
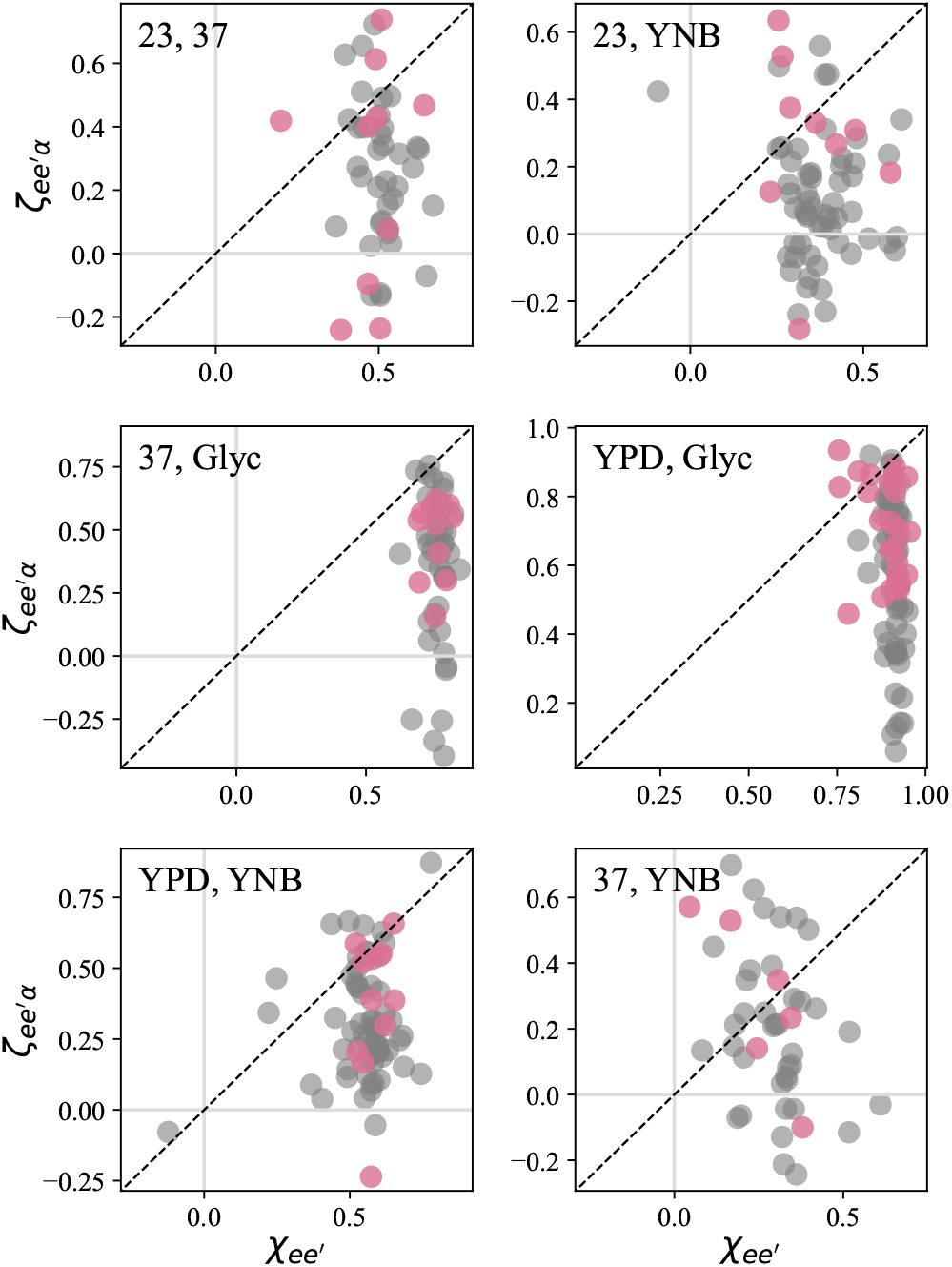
Test of the fully random pleiotropy model. Each panel shows the between-environment background correlation coefficient 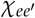 and focal mutation correlation coefficient 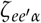 for each of the 87 focal mutations in each of six representative environment-pairs (see Fig. S8 for all other environment-pairs). Pink dots represent focal mutations which are in the WRE and SP limits for that environment-pair; gray dots represent focal mutations that are not. Note that 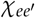 values for each mutation can be different due to missing data (mutational effects on some backgrounds are excluded due to experimental details, see Materials and Methods).

### Criss-cross environment predictions

#### Model

Our result in Eq. (6) provides a way to predict the slope of the relationship 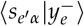 from measurements of the inverse relationship 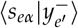 and the variance in background fitnesses in environments *e* and *e*^′^. We refer to this as a “criss-cross” environment prediction. Specifically, because 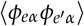 is symmetric with respect to the two environments, we see from Eq. (6) that the slope of the relationship 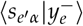 is given by the slope of the relationship 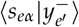 times the ratio of the variance in background fitness in environment *e*^′^ to the variance in background fitness in environment *e*. This provides a way to predict how the fitness effect of a focal mutation in environment *e*^′^ depends on the background fitness in environment *e*, without ever measuring its effect in environment *e*^′^.

This criss-cross prediction is exact in Eq. (6), and we therefore expect it to hold whenever the WRE and SP limits apply. However, we expect that it will often hold more generally, even when the SP limit does not apply. To see this, we note that in Eq. (5), the numerator of the relationship between *s*_*eα*_ and 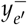 is given by 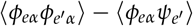. While the second term (which vanishes in the SP limit) is not symmetric, we expect it to be approximately symmetric whenever the correlations between epistatic terms involving the focal locus are similar to those between epistatic terms involving background loci (i.e. when the random pleiotropy approximation applies).

Because of this more general symmetry, we expect that focal loci need only satisfy either the random pleiotropy or the SP approximation for the criss-cross prediction to be valid. We can contrast this prediction with an alternative null expectation that the effects of mutations can be predicted from the general relationship between two environments (i.e. that the relationship between 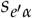 and *s*_*eα*_ for each focal mutation is the same as the relationship between 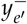 and 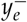).

#### Experimental Test

To test the criss-cross environment prediction, we use three measured components in the data for each environment pair (the fitness effect of the focal mutation in one environment, *s*_*eα*_, along with the background fitnesses in both environments, 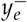 and 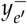), to predict the fourth (the fitness effect of the focal mutation in the other environment, 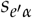).

As a baseline to compare against our generic null models, a natural hypothesis is that 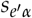 should be correlated with *s*_*eα*_ in the same way that 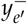 is correlated with 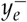. Specifically, this hypothesis predicts that the correlation between 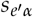 and *s*_*eα*_ is given by

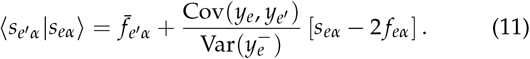

To test this, we fit the terms on the right-hand side of Eq. (11) from 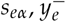, and 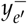, and use the result to predict 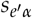 (Fig. 2A). As is apparent from the figure, this prediction performs poorly for most environment-pairs.

**Figure 2:**
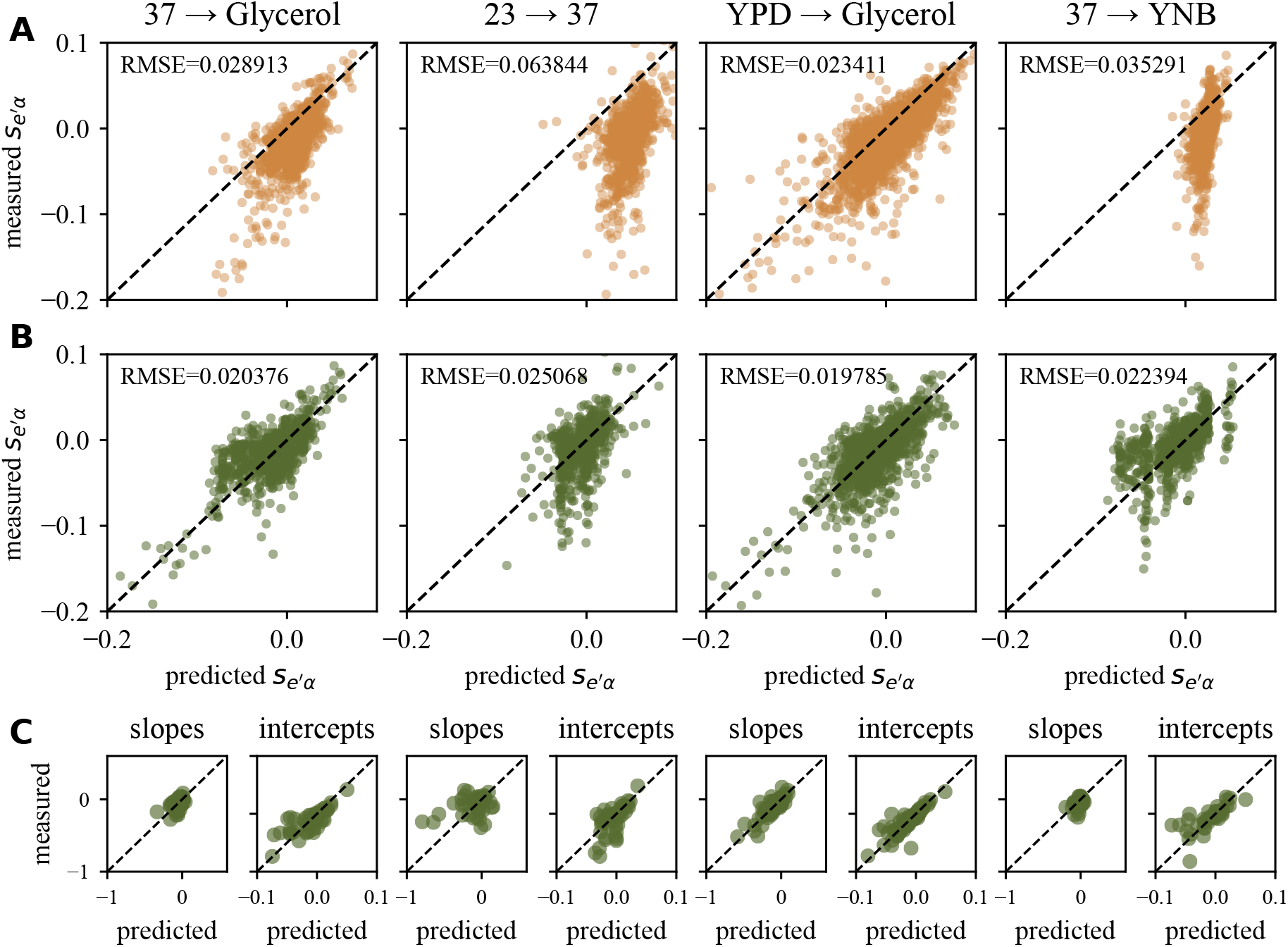
Test of the criss-cross environment prediction. Information from measurements of 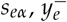, and 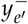 is used to predict 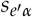. (**A**) Test of the baseline prediction that the correlation between fitness effects of focal mutations is identical to the correlation between background fitnesses for four representative environment pairs (see Fig. S9 for results for all other environment pairs). (**B**) Test of the criss-cross environmental prediction from our generic null model for the same four representative environment pairs. In panels **A** and **B**, each point corresponds to the predicted versus measured fitness effect of one focal mutation in one background genotype. (**C**) Predicted slopes and intercepts of the relationship between 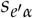 and 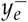 from our generic null model. Each point corresponds to the predicted versus measured slope or intercept for a given focal mutation.

We next test the criss-cross prediction of our generic null model, Eq. (6). Specifically, this predicts that the correlation between 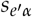 and 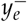 is given by

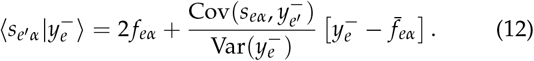

Here we use the average effect of the mutation across background fitness values in one environment as a prediction for the background effect in the other environment, 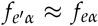. This is empirically supported between all environment combinations in our data. To test this prediction, we fit the terms on the right-hand side of Eq. (12) from the three measured values 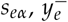, and 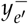, and use the result to predict 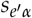 without using any information from this measurement (Fig. 2B). We see that this prediction performs much better than the baseline in Fig. 2A.

To highlight the performance of this prediction for both the slope and intercept in Eq. (12), we show the predicted versus measured values of these quantities for each focal mutation in Fig. 2C. We note that we expect these predictions to be accurate for focal mutations and environment pairs where the WRE and SP limits apply. However, as described above, even when these limits do not apply we still expect the criss-cross predictions to hold provided that 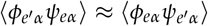. For most of our environment pairs, this is indeed the case, so our criss-cross predictions often hold even when the WRE and SP limits do not.

### Connectedness architecture

#### Model

We now turn to an analysis of global epistasis and pleiotropy in a multi-environment generalization of the “connectedness” model introduced by Reddy and Desai [2021]. In this model, each locus participates in some subset of *K* total “modules.” The output of each module is given by a connectedness model exactly as described by Reddy and Desai [2021]. The fitness across different environments is then determined by a weighted sum of these module outputs. Specifically, we have

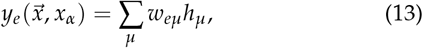

where the output *h*_*µ*_ of module *µ* is given by

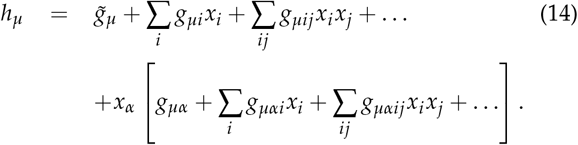

A key simplification of this model is that the module outputs *h*_*µ*_ depend on the genotype but not on the environment, which means the interactions among loci are fixed for all environments. In other words, the *g* terms do not depend on the environment; the environmental dependence is determined entirely by the *w*_*eµ*_ terms.

Consistent with the definitions introduced in Reddy and Desai [2021], we assume that there is a probability *ν*_*µi*_ that locus *i* participates in module *µ*. If it does, we assume the corresponding *g*_*µ*_ terms are independent and normally distributed with mean 0 and variance 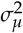. If it does not, the corresponding *g*_*µ*_ terms involving this locus are identically 0 (see Reddy and Desai [2021] for an alternative description of this model in terms of pathways within each module). These assumptions imply that

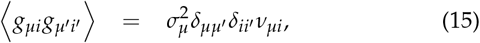

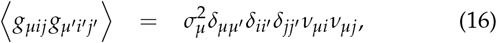

where the averages are over the distribution of the *g* terms. Analogous expressions hold for higher-order epistatic terms and terms involving focal loci such as 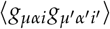. In addition, we assume that different epistatic terms are independent, so that expectations of products of different terms (e.g. 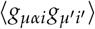) also vanish.

Without loss of generality, we will assume that 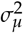 is tuned such that the total variance in the output of each module across all possible genotypes is 1 (we absorb differences in the importance of each module in the *w*_*eµ*_ terms instead). This implies that

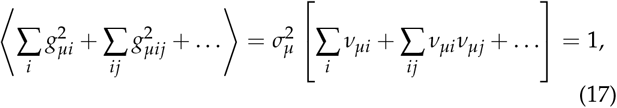

where all interaction coefficients are included in the expectation even those involving focal loci.

Given these assumptions, it is straightforward to calculate the quantities defined in Eq. (8). We find

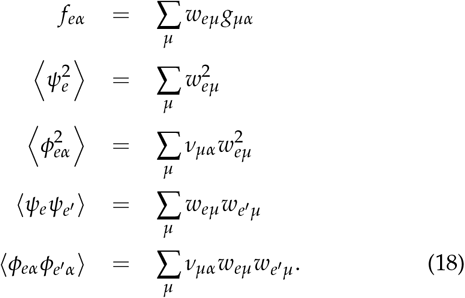

Here *f*_*eα*_ is the background averaged additive effect of mutation *α*. We can see from Eq. (18) that in the connectedness model there are a total of *FK* + *EK* + *K* parameters (*FK* parameters *g*_*µα*_, *EK* parameters *w*_*eµ*_ and *K* parameters *ν*_*µα*_) that determine the *E*^2^ + 2*E* total quantities. Provided that *K* < *E*, this represents a substantial simplification.

We can think of this connectedness model as representing a more general version of the fully random pleiotropy model. Specifically, the fully random model is similar in spirit to the connectedness model with only one module, and when we introduce multiple modules we allow for multiple ways in which environments can be correlated. Because fitness in each environment depends on the modules in different ways, the participation of different loci in different modules can then allow them to have distinct patterns of pleiotropy.

#### Experimental Test

To fit our empirical data to the connectedness model architecture, we aim to find the set of *FK* + *EK* + *K* connectedness model parameters (i.e. the *FK* parameters *g*_*µα*_, *EK* parameters *w*_*eµ*_ and *K* parameters *ν*_*µα*_) that best fit the data (i.e. the *FE* terms *f*_*eα*_ along with the *E*^2^ + 2*E* variances and covariances 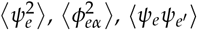, and 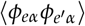) using the relationships described in Eq. (18). To do so, we minimize the following loss function,

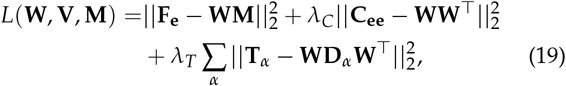

using the procedure described in the Supplementary Text. The matrix **F**_**e**_ consists of 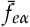 terms for each locus in each environment, **C**_**ee**_ and **T**_*α*_ are covariance matrices of *ψ*_*e*_ values and *ϕ*_*eα*_ values, respectively, **W** has matrix elements *w*_*eµ*_, **M** has elements *g*_*µα*_, **D**_*α*_ = diag(**V**_*α*_), and **V** contains *ν*_*µα*_ values for each locus, for all modules. Specifically, **F**_**e**_ ∈ ℝ^*E*×*F*^, **W** ∈ ℝ^*E*×*K*^, **M** ∈ ℝ^*K*×*F*^, **C**_**ee**_ ∈ ℝ^*E*×*E*^, **T**_*α*_ ∈ ℝ^*E*×*E*^ for *α* = 1,.., *F* and **V**∈ ℝ^*F*×*K*^. The first term is reminiscent of the SSD framework in Petti et al. [2023] used to fit isolated additive effects, the second term describes the background variance between environments, and the last term describes the epistatic contributions of each locus. Using a train, validation, test procedure the best-fit hyper-parameters are *λ*_*C*_ = 0.01, *λ*_*T*_ = 10^6^, and *K* = 4 (see Supplementary Text).

We then analyze how well this connectedness model predicts the dependence of the fitness effect of each focal mutation in each environment on the background fitness in each other environment. This is done by comparing the slopes (Fig. 3A) and intercepts (Fig. 3B) of the dependence (as defined in Eq. (6)) predicted from the model to those measured from our data. We see that the connectedness model provides a good fit for focal loci and environment-pairs which fall into the SP limit, suggesting that where this assumption holds, the model does provide a good description of lower-dimensional structure in the data.

**Figure 3:**
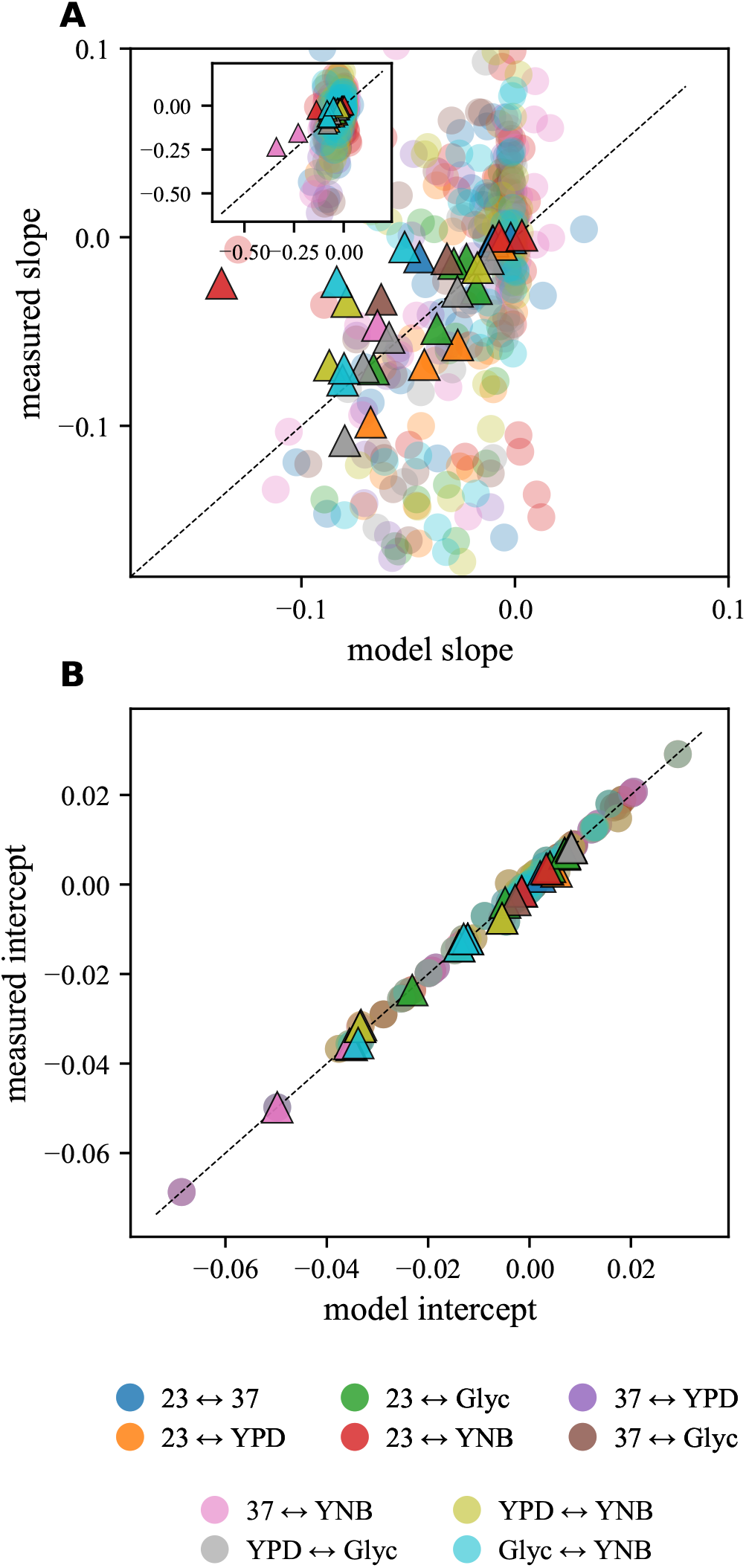
Test of the connectedness model. **(A)** Observed versus predicted slope of the dependence of the fitness effect of each focal mutation in each environment on the background fitness in each other environment, 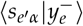. Triangles denote focal mutations and environment-pairs for which the SP limit holds, while circles denote those where it does not. **(B)** Observed versus predicted intercept of the dependence of the fitness effect of each focal mutation in each environment on the background fitness in each other environment, 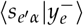.

### Evolutionary tradeoffs borne out by interaction structure

In the previous section, we found that the data is well described by a lower-dimensional framework: environments that depend differently on a set of modules, and loci with varying degrees of participation in each module. Here we ask whether the interaction structure of this low-dimensional framework implies different evolutionary outcomes (specifically, if tradeoffs between fitness across environments can result from these generic models rather than specific biological mechanisms). We introduce three models with different assumptions about how loci participate in modules and analyze the long term effects of adaptation given different environment–module dependencies.

The three models we analyze are (1) *independent modules* where the participation of a locus is drawn independently for each module, (2) *isolated modules* where loci only participate in one module, and (3) *homogeneous modules* where loci have the same participation in all modules. Structures for each of these models are illustrated in Fig. 4A. The interaction structure observed in the data is described in Fig. 4B, and the distributions of participation for each locus (*ν*_*µα*_) are analyzed in Fig. 4C. We find loci in this dataset are mostly described by the independent modules model, with some loci having isolated effects on one of the four modules (Fig. 4C, third column).

**Figure 4:**
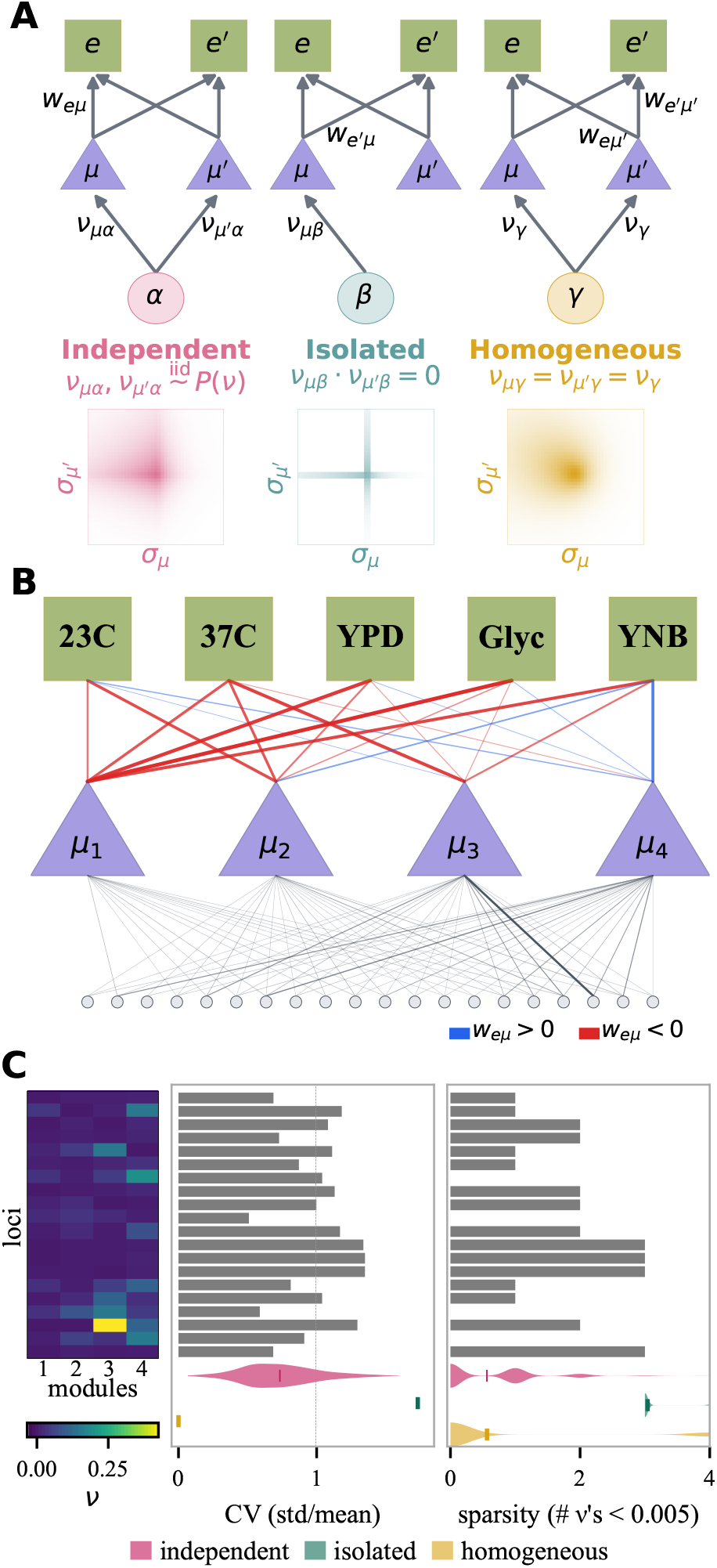
Structure of locus participation in modules. **(A)** The three models studied and their associated JDFE. Loci are either all *independent* (*ν*_*µα*_ and *ν*_*µα*_ sampled iid from *P*(*ν*)), *isolated* (only participating in one module), or *homogeneous* (same level of participation in all modules). Module outputs *h*_*µ*_ = 10 and *h*_*µ*_′ = −5 are used for the JDFE of each model. **(B)** Empirical genotype–phenotype map resulting from the minimization of Eq. (19). Thickness and opacity of each line corresponds to the weighting of each module by each environment (blue = positive dependence and red = negative dependence) or the participation of loci in each module. **(C)** Given **A** and **B**, which model structure do the empirical loci follow? The value of *ν*_*µα*_ for each locus/module appears across rows/columns in the heatmap. The coefficient of variation, CV (standard deviation / mean) and sparsity (# of *ν*_*α*_ values < 0.005) are plotted for each locus compared to loci sampled from the JDFE of each model.

Motivated by this simplification, in this section we analyze the implications of this generic null model of complex traits for the outcomes of long-term evolution in fluctuating environments. To do so, we first generalize the connectedness architecture introduced above, removing the discrimination between focal and non-focal mutations. We then derive the joint distribution of fitness effects (JDFE), the probability that a new mutation has fitness effect 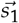 across these environments, in these models in the WRE and SP limits, under three different classes of assumptions about how loci participate in different modules. Finally, we use these JDFEs to ask what happens to a population evolving under selection that alternates between two environments. Specifically, we calculate whether such a population is expected to reach a stable steady-state fitness (and if so what tradeoff between environments this fitness represents), and how this depends on both the modular architecture and the relationship between the two environments.

#### JDFE in the connectedness model

To calculate the JDFE of an arbitrary new mutation, we move away from the simple experimental scenario described above, and consider a version of the connectedness model that does not distinguish between focal and background loci. Specifically, we write the output of module *µ* as

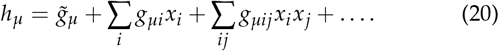

The fitness of genotype 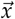 in environment *e* is then given by

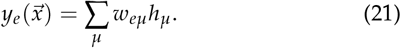

Without loss of generality, we rescale the *g* and *w* terms such that 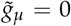 for all 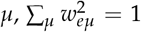, and 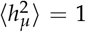 (and hence the variance in fitness in each environment is 1).

As above, we assume that the output of each module *µ* is determined by a connectedness model architecture, in which locus *i* participates in a fraction *ν*_*iµ*_ of the pathways in module *m*. In this framework, Reddy and Desai [2021] show that the effect, *σ*_*iµ*_, of locus *i* on the output of module *µ* is

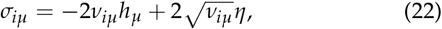

where *η* is normally distributed (with mean 0 and variance 1) across background genotypes at other loci. This implies that the distribution of effects on the output of this module is given by

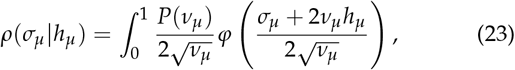

where *h*_*µ*_ is the current output of module *µ, φ* is the standard normal pdf, and *P*(*ν*_*µ*_) is the probability a locus in module *µ* has a participation fraction *ν*_*µ*_ (i.e. the distribution of variance fractions, DVF, as defined by Reddy and Desai [2021]). We will assume that this DVF is exponential, 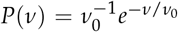; in this case we can integrate Eq. (23) to find

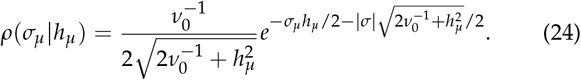

In other words, the distribution of module effects follows a Laplace distribution when the DVF is exponential.

To compute the JDFE, we must generalize this result to analyze the joint distribution of effects on the outputs of multiple modules. Because the joint distribution of outputs of multiple modules is multivariate normal in the WRE and SP limits, for the case of *n* modules we have

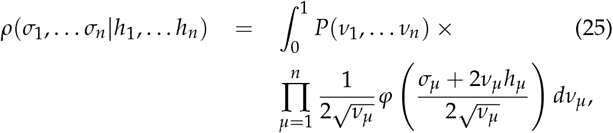

where *P*(*ν*_1_, *ν*_2_, … *ν*_*n*_) is now a joint distribution of variance fractions (JDVF) across the *n* modules. The JDFE follows straightforwardly from Eq. (25); for each environment *e* we simply weight the output of each module with the corresponding *w*_*eµ*_.

Our result in Eq. (25) and the corresponding JDFEs depend on the joint distribution of variance fractions. We consider three different models for this JDVF. In the first model, we assume the participation of loci in one module is independent from their participation in others. In this *independent modules* case, we have 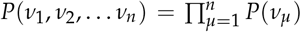, and the joint distribution of module outputs becomes

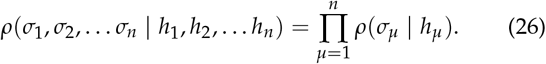

In the second model, we assume that each locus participates in only one module. In this *isolated modules* case, we have 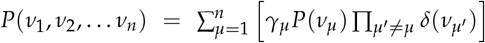, where *γ*_*µ*_ is the fraction of sites that participate in module *µ*. The joint distribution of module outputs is then

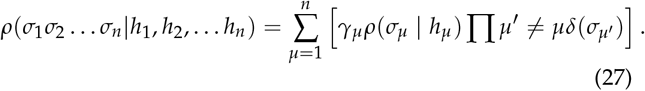

Finally, in the third model, we assume that a given locus participates equally in all modules. In this *homogeneous module* case, we have 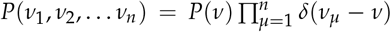, and the joint distribution of module outputs becomes

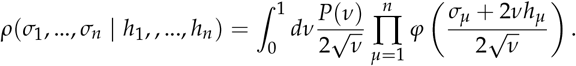

In this case, using the exponential *P*(*ν*) from above, we find

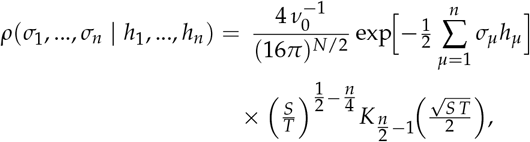

where 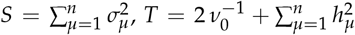 and *K*_*l*_ is the modified Bessel function of the second kind.

#### Long-term evolution in fluctuating conditions

Long-term adaptation in a fluctuating environment is driven by the fitness effects of new mutations across environmental conditions the population has or will experience, along with the corresponding probabilities that these mutations fix or are driven to extinction. We imagine our starting population has fitness 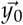 across some set of environments (with each component of this fitness vector representing the fitness in one of the environments). The JDFE, 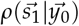, dictates the probability that a new mutation has fitness effect 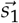 across these environments. Depending on which environment the population is currently experiencing, the fixation probability of this new mutation will then depend on the corresponding component of 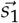. If this mutation fixes, the population is now at some new fitness, 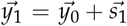. The joint distribution of fitness effects now shifts to 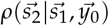, and the process continues (with selection now potentially acting on fitness in the other environment).

In general, after a set of *l* mutational steps, the JDFE for the next step is some complicated function of all previous steps, 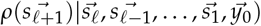. In principle, this dependence of the JDFE at each step on all previous mutations can lead to a complex history dependence in evolutionary trajectories. However, as described by Reddy and Desai [2021], in the connectedness model this history dependence vanishes, and the JDFE depends only on the current fitness. In other words, 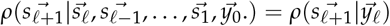.

Given this history independence and each JDFE calculated above, we can now analyze expected long-term outcomes of evolution given some specified dependence of the fitness in different environments on these module outputs and timecourse of fluctuating environmental conditions. For simplicity, we will focus on a two-environment and two-module system, in which the fitness in environment *e* and *e*^′^ can be written as

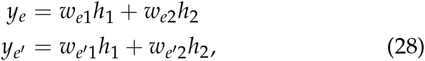

where *w*_*e*1_ and *w*_*e*2_ represent how the fitness in environment *e* depends on the output of modules 1 and 2 respectively, and analogously 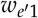 and 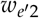 represent how the fitness in environment *e*^′^ depends on these modules. We can visualize the vector 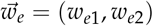 as pointing in the direction that selection favors in the space of module outputs (Fig. 5A).

**Figure 5:**
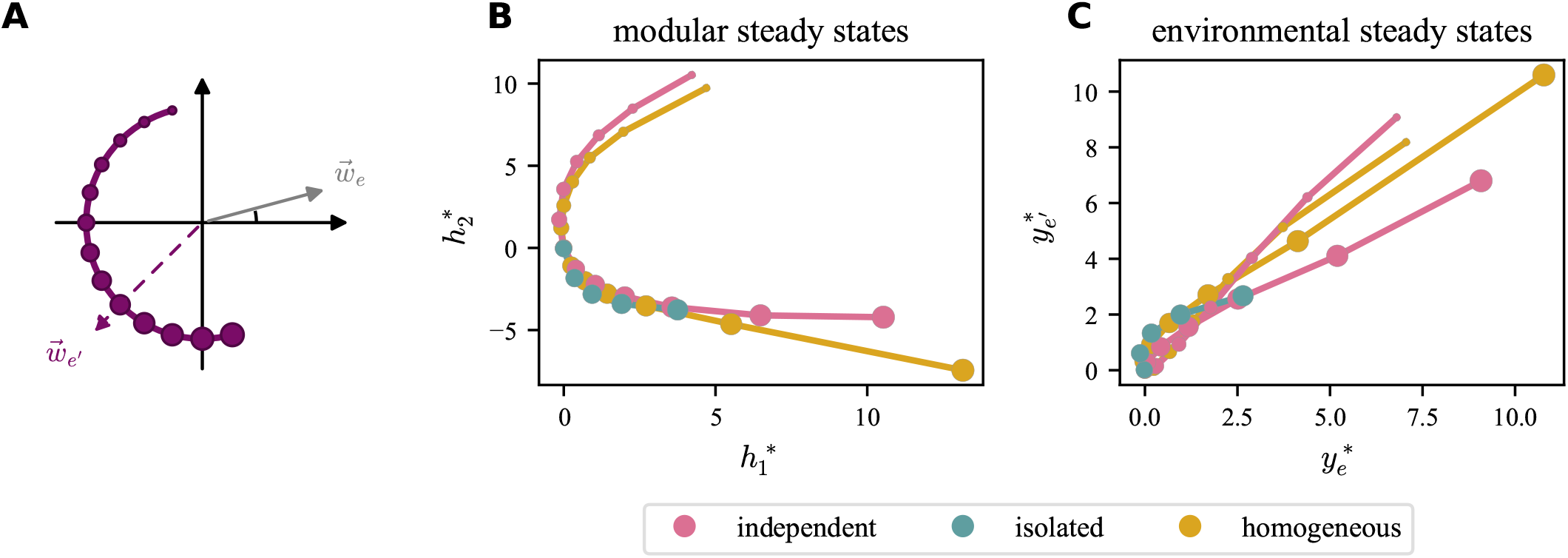
Long-term evolutionary steady states. **(A)** We can visualize the effects of module outputs on fitness in the two environments as vectors 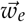 and 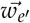. Here we show a case where 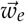 is at 15° and 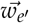 ranges from 105° to 285°. **(B)** Outputs of modules 1 and 2 as a function of the angle of vector 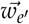. Results for each JDVF case are shown in different colors. **(C)** Steady state fitness reached in environments *e* and *e*^′^. Marker size indicates the angle of selection in environment *e*^′^, 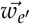.

We now imagine that evolution proceeds through a series of mutational steps. For concreteness, we imagine that the population experiences a fluctuating environment that shifts at each mutational step. We assume that at each step the JDFE is given by the expressions above, and that the fixation probability of a mutation is given by twice the selective effect of the mutation in that environment (as expected in the strong selection weak mutation limit, with no clonal interference). Note that this means that if we are currently in environment *e*, a mutation which has an effect 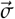 on the module outputs has fixation probability 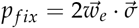.

We expect long-term evolution in this model to result in a steady state fitness if it leads to a state where selection in the two environments on average moves the module outputs in equal and opposite directions. In other words, we expect a steady state at the modular output values 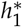 and 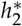 for which 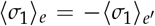 and 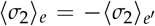, where ⟨*σ*_*i a*_⟩ is defined as the average effect of a fixed mutation in environment *a* on module *i*. We can compute these average effects using the JDFEs and fixation probabilities as described above. We find that whether such a steady state exists (and the nature of the tradeoff it implies when it does) depends on the JDVF and the dependence of fitness on the module outputs in the two environments, 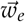 and 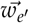.

We begin by considering the isolated modules case, in which each mutation affects only one of the two modules. Here, we find that if *w*_*e*1_ and 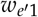 are the same sign, there is no tension between the two environments, and evolution can favor ever-increasing (or ever-decreasing) the output of module 1. Similarly, if *w*_*e*2_ and 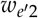 are the same sign, evolution can favor ever-increasing or ever-decreasing the output of module 2. In either of these cases, long-term evolution will not result in a steady state. However, if *both* modules are individually under opposing tension, (i.e. *w*_*eµ*_ and 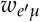 have opposite signs for both *µ* = 1 and *µ* = 2), then a tradeoff exists and long-term evolution leads to a steady state, with steady state values of module outputs and fitness in each environment depending on the relative orientations and magnitudes of 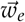 and 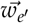, as illustrated in Fig. 5B and Fig. 5C respectively.

In the independent modules and homogeneous modules cases, we find that the steady state exists provided only that 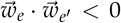. This weaker condition is sufficient in these cases since each locus participates in multiple modules with differing (independent case) or correlated (homogeneous case) effects, allowing tradeoffs to exist at the locus level even when selection acts in the same direction on the output of a single module. In this case, selection in the two environments tends to move module outputs overall in opposing directions, so that no combination of modules can be improved indefinitely and evolution will eventually reach a steady state, as illustrated in Fig. 5B,C.

Broadly speaking, we see from Fig. 5 that regardless of which JDVF case we consider, the steady state module outputs are close to 0 when the two environments select on them in nearly opposite directions 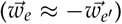, and the corresponding fitness in both environments is low. As selection on the two modules becomes less directly opposed, we instead reach a steady state with nonzero module outputs (which reflects the average tension between the selection pressures on each module) and the steady state fitnesses in both environments increase. The quantitative details of this dependence depends on the assumptions underlying the JDVF.

We note that in this analysis, we have focused on the connectedness model, where history-dependent effects can be neglected. In other architectures (which may still satisfy the WRE and SP approximations), this history dependence may be important and could alter the steady state. In addition, we have implicitly assumed in this analysis that the genome is effectively infinite, so that mutations with all possible effects on module outputs are always available regardless of the current genotype. A finite genome would induce history-dependent effects and fluctuations around these averages, which could also lead to changes in the steady state.

Finally, we have focused on a simple two-environment example in this section. It is straightforward in principle to extend this to higher dimensions, and the existence and properties of long-term evolutionary steady states can be calculated using the same logic as described above. However, assumptions about the order and timing of environmental shifts, as well as of the tensions (or lack thereof) between module outputs across many environments, will play an important role.

## Discussion

A large body of experimental and theoretical work has demonstrated that epistasis is widespread and often leads to predictable global patterns such as diminishing returns and declining adaptability across microbial and viral, systems [Kryazhimskiy et al., 2014, Couce and Tenaillon, 2015, Lenski et al., 2015, Good and Desai, 2015, Perfeito et al., 2014, Bakerlee et al., 2022, Lyons et al., 2020, DelaFuente et al., 2024, Bourg et al., 2024, González-González et al., 2024, Witte et al., 2023, Moulana et al., 2022]. Other studies have shown that mutational effects vary strongly across environments and phenotypes, revealing both persistent pleiotropy and local modularity [Martin and Lenormand, 2015, Geiler-Samerotte et al., 2020, Kinsler et al., 2020, Wang et al., 2024, Jerison et al., 2017]. Work on fluctuating and changing environments has further emphasized that tradeoffs, modularity, evolvability, and even the meaning of fitness itself depend on environmental structure and temporal variation [Kashtan et al., 2007, Tikhonov et al., 2020, Sæther and Engen, 2015, Abreu et al., 2024].

However, much less has been understood about how epistasis and pleiotropy interact — specifically, how the background dependence of mutations changes across environments. This study addresses that gap by extending generic models of global epistasis into a multi-environment setting and showing that many cross-environment patterns can emerge from widespread random interactions combined with low-dimensional shared structure. Using mutation-effect measurements across many yeast genetic backgrounds and environments, we find that fully random pleiotropy is insufficient to explain the data, but that a connectedness/modular model captures much of the observed covariance structure for the subset of mutations which exhibit structured pleiotropy. The existence of a lower-dimensional framework encompassing a shared interaction structure between environments is not *a priori* expected to represent the data. However, our finding that this modular structure is representative is evidence that there is embedded information about mutations between all measured environments due to this shared structure. We show this information can be identified through the cross-environment global epsitatic relationship. In this way, our work connects analysis of global epistasis, pleiotropy, modularity, and fluctuating-environment evolution by proposing that many tradeoffs and patterns of cross-environment predictability arise from shared latent interaction structure rather than highly specific mechanistic constraints.

A particularly close point of comparison is the recent work of Ardell et al. [2024], which showed that distributions of mutational effects across environments can appear remarkably environment-independent despite extensive microscopic epistasis. Ardell et al. [2024] argued that global epistatic structure is largely shared across environments and differs primarily through an environment-specific “pivot” growth rate, implying that adaptation in one environment predictably constrains adaptation in others. Our work extends and generalizes this perspective by explicitly analyzing the covariance structure of mutation-specific epistatic interactions across environments. Rather than assuming a largely invariant global structure, we ask when shared cross-environment interaction geometry emerges and when it breaks down. We find that while some mutations satisfy a strong structured pleiotropy (SP) limit consistent with shared latent interaction structure, many mutations do not, indicating that cross-environment predictability is only partial and mutation-dependent. In addition, our connectedness model provides a more general framework in which environments can weight shared latent modules differently, allowing both positively and negatively correlated pleiotropic effects to emerge naturally.

A key limitation of our work is that most mutations do not satisfy the stronger structured pleiotropy (SP) limit, suggesting that shared cross-environment interaction structure is only partial and may not fully capture the diversity of mutational effects. In addition, our analysis of long-term evolutionary steady states relies on highly idealized assumptions, including a strong-selection weak-mutation (SSWM) regime without clonal interference, an effectively infinite genome where beneficial mutations are never exhausted, and the absence of ecological feedback between populations and environments. The framework also does not explicitly model evolutionary history dependence, so its conclusions about long-term adaptation and tradeoffs remain theoretical extrapolations from single steps in each environment. In addition to these assumptions, we also do not explore the regime of a large number of environments or modules. Further analysis using our framework in this regime could probe more realistic evolutionary dynamics.

Another important direction for future work will be determining how much of the observed cross-environment structure reflects truly shared latent physiology versus statistical correlations induced by background fitness structure. Measuring a much larger and more diverse set of mutations across the same genetic backgrounds could help reveal additional latent modules and clarify whether mutations outside the SP limit are genuinely environment-specific or simply weakly coupled to shared interaction structure. On the theoretical side, future work could investigate how the dimensionality and overlap of latent modules shape long-term evolutionary dynamics in fluctuating environments, how cross-environment epistatic structure evolves over time, and whether quantities such as environmental memory, adaptability, or tradeoff strength can be predicted directly from the covariance structure of mutational interactions. More broadly, extending these models beyond static pairwise environments to temporally fluctuating or spatially heterogeneous settings may help unify ideas from global epistasis, modularity, and fluctuating-environment evolution within a common statistical framework.

## Code and Data Availability

All code and data presented in this manuscript are available at https://github.com/caelan-brooks/ep_pleio_int. Sequencing data is available on SRA (accession number PRJNA1536643).

## Materials and Methods

### Strain construction

The yeast strains used in this study are derived from Johnson et al. [2019]. Our strain library consists of 87 of the 176 background genotypes used in this earlier study (those referred to as E2); strains excluded after analysis by Johnson et al. [2019] were also excluded here. These 87 background genotypes are haploid F1 segregants generated from a cross between a laboratory strain (BY) and wine strain (RM), and differ at ~42,000 background loci. In each of these background strains, Johnson et al. [2019] introduced a total of 96 gene disruption mutants using a barcoded version of a diverse Tn7 plasmid library created by Kumar et al. [2004]. Of these gene disruption mutants, 91 were chosen from a larger set to be those with significant fitness effects in our rich laboratory environment (YPD at 30C), and 5 were chosen as putatively neutral controls. This created a total library of 8352 strains (96 focal gene disruption mutations in each of 87 background genotypes). Each strain contains redundant barcodes, with each barcode mapping to a specific background genotype and focal mutation. For further details of strain construction and genotypes, see Johnson et al. [2019].

### Bulk Fitness Assays

Our trait of interest lends itself especially well to high-throughput phenotypic measurements. Relative growth rate of each individual mutation, which we call ‘fitness’, is measured through sequencing assays. We perform barcode based assays in bulk, pooling all mutations against a single segregant background, and read out relative frequencies by amplicon sequencing a barcode region. This all-to-all competition necessarily assumes no frequency dependent selection or interactions (except competitive exclusion) of any kind.

All bulk fitness, growth assays were performed in unshaken, flat-bottomed, polypropylene 96-well plates, using a liquid-handling BiomekFX robot (Beckman Coulter). Fitness assays were conducted in 5 different growing conditions, as outlined in Table S1. All segregant libraries were revived from frozen stock, and we performed a 1 : 2^6^ dilution in 128 µlat each transfer, allowing for 6 generations of growth per day over 8 days. At 7 different time points, a sample of the full saturated culture was taken, and pelleted for downstream processing. Details can be found in SI: Material, Methods and Protocols

All cell pellets were processed to extract genomic DNA from a protocol outlined in Nguyen Ba et al. [2019], using silica columns and cell lysis with zymolyase. Any genomic DNA not used for library preparation was stored at −20°C. Since time point zero is grow-up to allow cells to acclimatize to each environment, we do not sequence that or include it in our analysis. We sampled all time points from generation 6 of the fitness assay to generation 42. For our sequencing library, we produced Illumina compatible, dual indexed, UMI (Unique Molecular Index) tagged amplicons using a 2 step PCR protocol. The protocol follows principles from Levy et al. [2015] and uses primer sequences from Johnson et al. [2019]. The same pool of fragments was used in both lanes, however due to other experimental considerations, significantly different total reads per sequencing lane were obtained. Details can be found in SI: Material, Methods and Protocols

### Inferring mutational fitness from sequencing data

We infer fitness from changing barcode frequencies over the fitness assay. To go from FASTQ files generated by the sequencer to barcode counts, and eventually frequencies, we use the methods and code outlined in Ba et al. [2022], using only the forward read from each sequencing run. Each read is processed by removing possible sequencing artifacts and parsing using regular expression (regex) programming in python. We look for a 16 base pair region before the barcode (with up to 1 error allowed) and the 40 base pair region after the barcode (upto 3 errors allowed). The barcode regions follow a specific pattern of NNNN-TG-NNNN-TG-NNNN-TG-NNNN-TG-NNNN; which is also built into the regex using lazy matching. Each barcode sequence is then combined with the UMI’s (designed to be the first 7 base pairs of the read). We typically find that around 90% of our sequence data is usable after this parsing, giving us putative barcodes. These putative barcodes follow this defined structure, but may still have some errors. Since we know the list of true barcodes that must be present in the experiment, we can correct against the the list using error correction algorithm from Ba et al. [2022], allowing up to 3 mismatches in a 28 base pair barcode. We expect errors to be rare and can use direct lookup of errors to the true dictionary, and correct as many reads as possible. Finally, we remove any duplicate reads that share the same barcode-UMI combination (any repeats are likely being generated due to PCR amplification biases), counting barcodes occurrences at each time point, and matching counts from different time points to get barcode trajectories. The complete sequencing data used in our analysis comes from 2 different lanes. Hence, this analysis was performed twice. To get final read counts, counts from each barcode and each time point of the lanes was summed up. A separate count was maintained for each barcode, and each mutation (edge) was redundantly barcoded. This resulted in each mutation (edge) having multiple independent barcode counts and frequency trajectories.

We inferred the fitness of each strain from the barcode counts using the methods and code from Johnson et al. [2019]. For time-points and replicates with enough data, we measure the log-frequency slope for each set of successive time points, scale by the fitness effect of known neutral mutations, and average across all time intervals in the assay. Since each mutation (“edge”) has been redundantly barcoded, we generate a group of frequencies for each with values that putatively measure the same fitness effect. Because of this redundancy, we can use a log-likelihood ratio test to exclude barcodes that differ from the median fitness of the group corresponding to the same mutation. All our fitness estimates are reported in per generation of growth.Details can be found in SI: Material, Methods and Protocols

### Background Fitness Estimation

We revive frozen stocks of the strain backgrounds and a florescent reference strain from Jerison et al. [2017], containing mCitrine. We grow up both types of strain to saturation, and then co-culture them for a 2 time point assay, separated by 6 generations (12 for YNB). We count the number of cells in the population of dark cells (strain backgrounds) and fluorescent cells (reference), using the change in these frequencies to estimate fitness. We use tools from the fcparser and cytoflow libraries to analyze this data, fitting a Gaussian Mixture model to separate the cells into the 2 types. Once counted, fitness against the reference florescent strain was measured by calculating 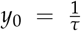 ln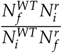 where *τ* = 6 and values in the natural log and final and initial counts of the reference and WT strains. For YNB, a 12 generation assay was conducted, and hence *τ*_*YNB*_ = 12.

## Acknowledgments

The authors would like to thank members of the Desai lab for helpful conversations and feedback, as well as members of the Harvard Bauer Core facilities for help with sequencing. This work was supported by NIH grants R01-GM104239 and R35-GM161612. CB acknowledges funding from the NSF Graduate Research Fellowship Program under Grant No. DGE-2140743.

## Supplemental Material

### Supplementary Figures

**Figure S6:**
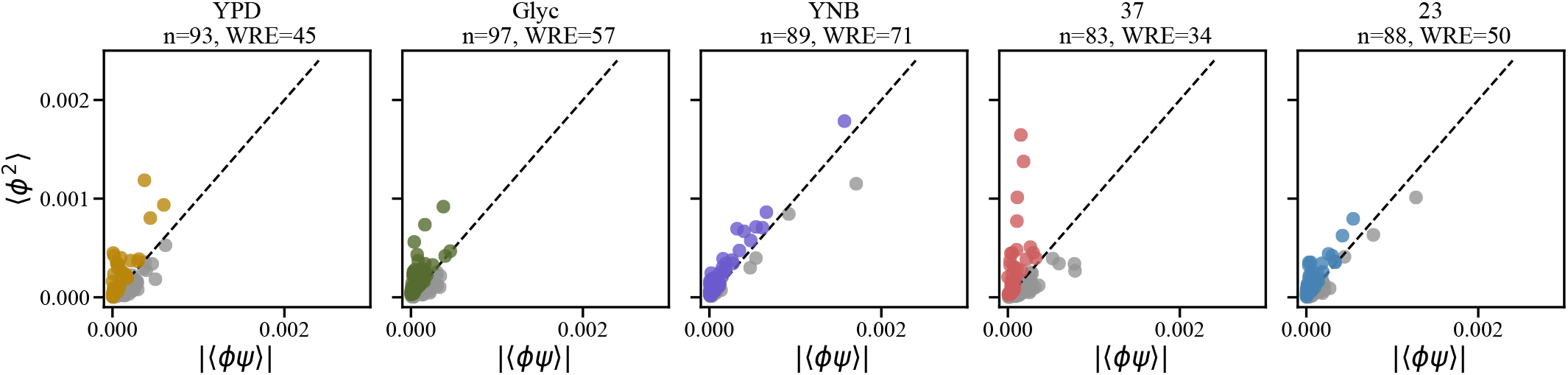
Applicability of the WRE approximation in each environment. Each point represents the values of 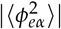 and |⟨*ϕ*_*eα*_*ψ*_*eα*_⟩| for one focal mutation in each of our five environments. Environment identities, the total number of focal mutations, *n*, and the total number of these focal mutations in the WRE limit in that environment (defined as those for which 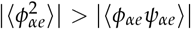 are shown at top of each panel.

**Figure S7:**
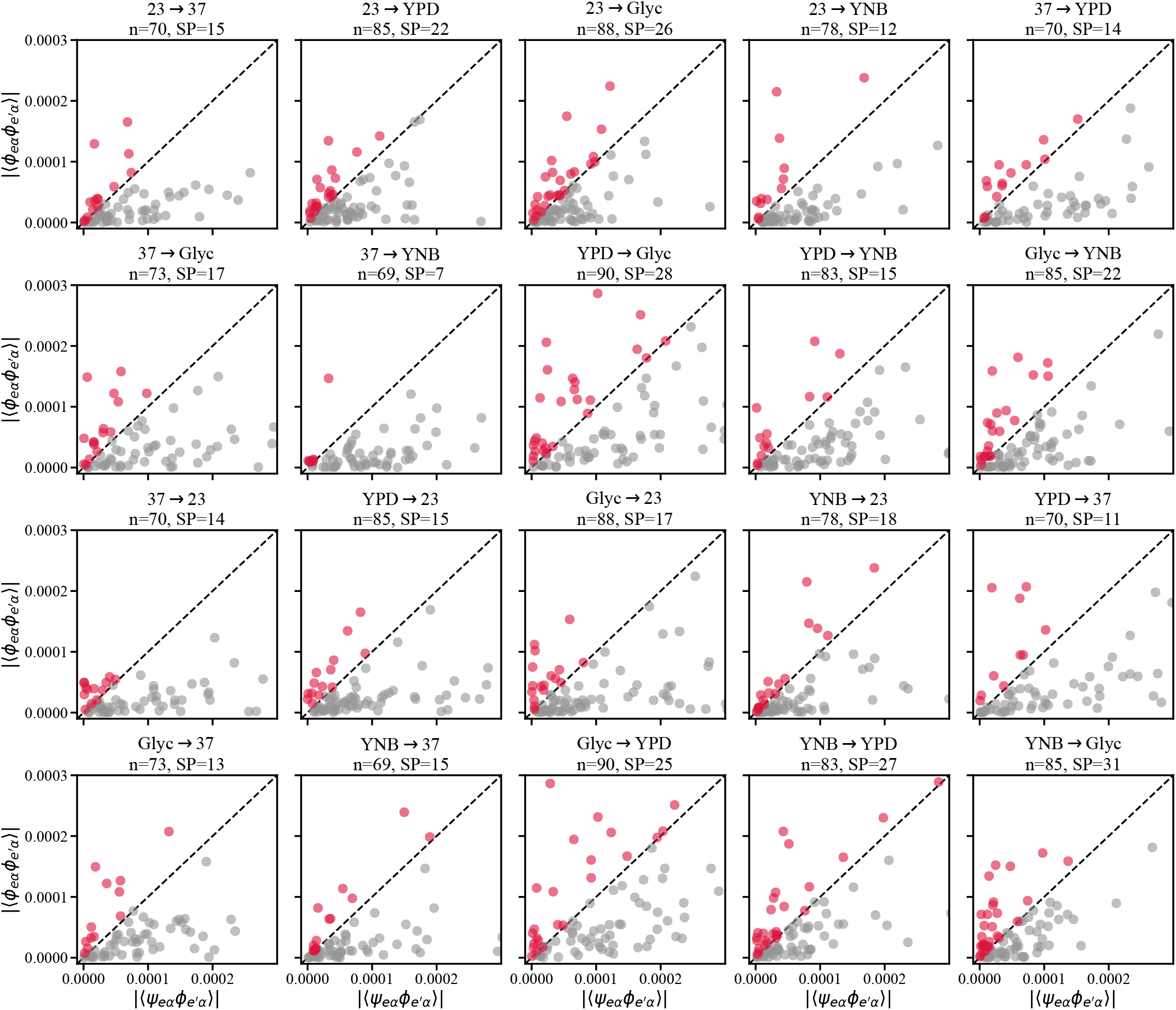
Applicability of the SP approximation for each environment-pair. Each point represents the values of 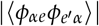 and 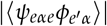 for one focal mutation in each environment-pair. Environment identities, the total number of focal mutations, *n*, and the total number of these focal mutations in the SP limit for that environment-pair (defined as those for which 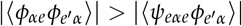) are shown at top of each panel.

**Figure S8:**
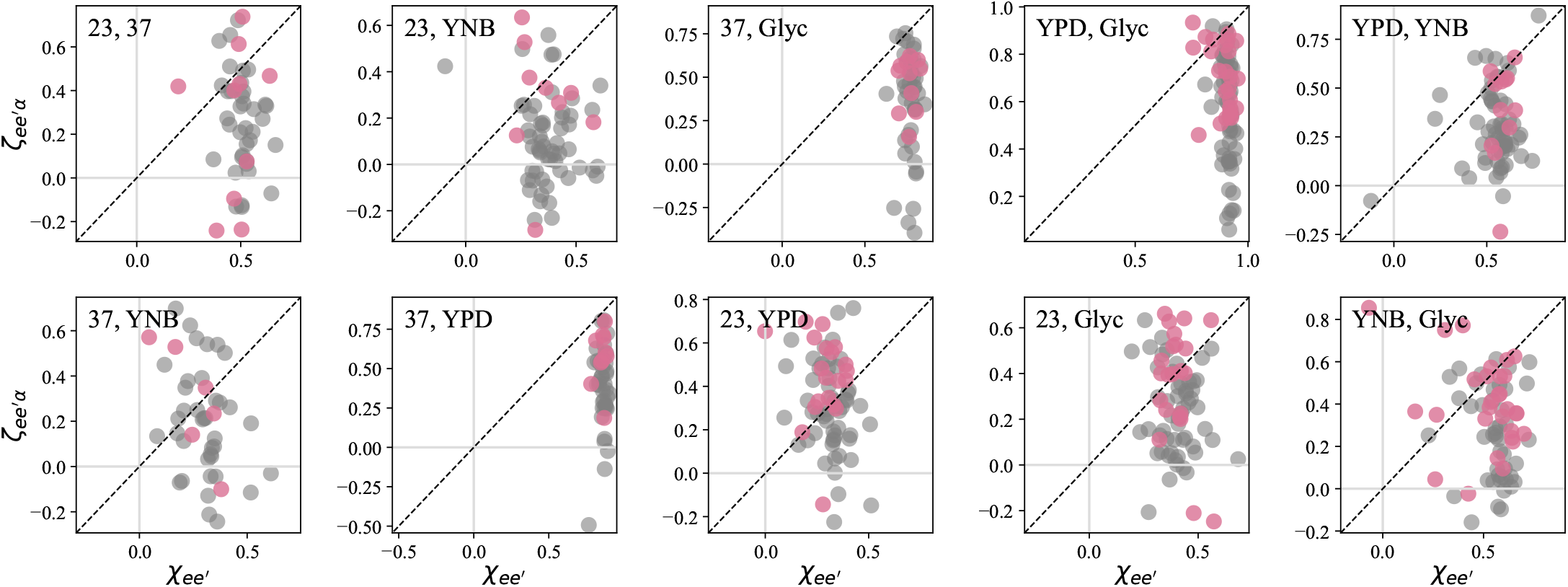
Test of the fully random pleiotropy model. Each point represents the correlation coefficient of the fitness effect of a given focal mutation across a given environment-pair, 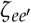, plotted as a function of the correlation coefficient of background fitness values in that environment-pair, 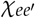. Focal mutations that are in the WRE and SP limits in a given environment-pair are shown in pink, while all others are shown in gray. Note that 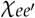 values for each focal mutation can be different due to missing data.

**Figure S9:**
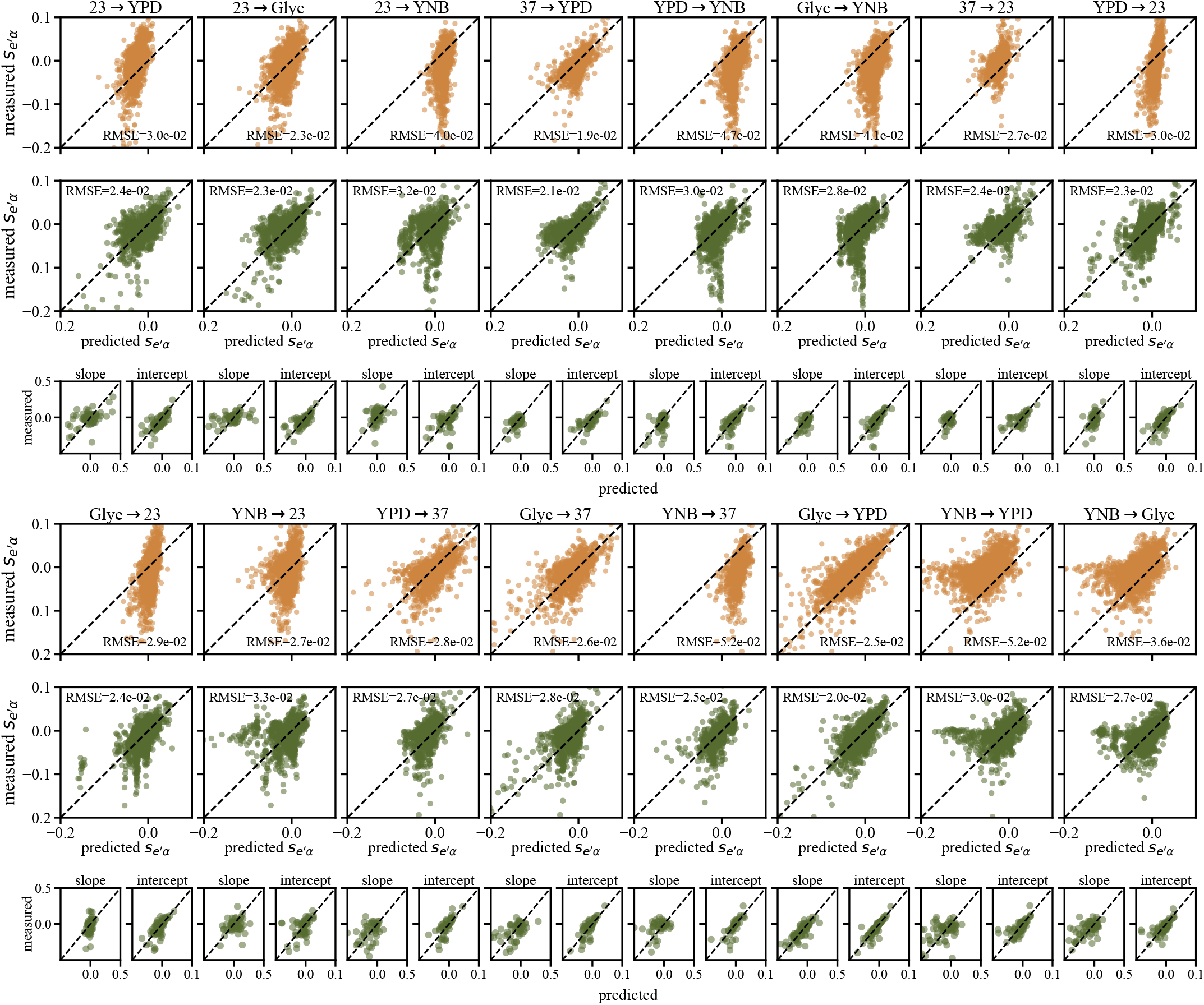
Test of the criss-cross environment prediction in all environment-pairs. Yellow plots show test of the baseline prediction that the relationship between fitness effects of focal mutations is identical to the relationship between background fitnesses. Large green plots show test of the criss-cross environmental prediction from our generic null model. Small green plots show predicted slopes and intercepts of the relationship between 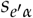 and *seα* from our generic null model.

### Supplementary Text

#### Generalization to multiple focal loci

Throughout the text we have focused on the case of a single focal locus (*F* = 1). However, the data analyzed includes focal mutations all made on the same set of background genotypes. We now consider how our results can be generalized to the case where we measure the fitness effect of multiple (*F* > 1) focal loci (as before, in *E* different environments across a large number of genetic backgrounds that vary across *L* background loci). Because we do not consider the effects of making multiple focal mutations simultaneously, we can treat each focal locus separately. In other words, for focal locus *γ*, we can write the fitness in environment *e* as

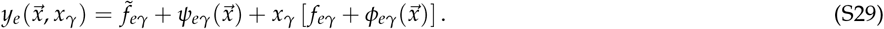

However, it is important to note that all terms in this equation now depend on which focal locus we are considering, and include contributions from terms involving all the other focal loci.

To see this, consider the case of two focal loci, which we refer to as *x*_*α*_ and *x*_*β*_. We have

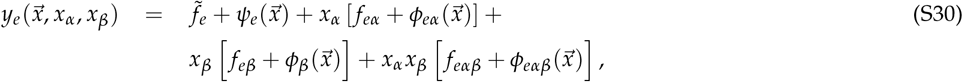

where *ψ*_*e*_, *ϕ*_*eα*_, and *ϕ*_*eβ*_ are as defined in Eq. (2) and Eq. (3) respectively, and *ϕ*_*eαβ*_ involves all *f* terms that depend on both focal loci and one or more background loci. We can immediately see that if we consider only focal locus *x*_*α*_ (and hence set *x*_*β*_ = −1 throughout), we can define 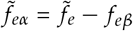 and *ψ*_*eα*_ = *ψ*_*e*_ − *ϕ*_*eβ*_, and redefine *f*_*eα*_ → *f*_*eα*_ *− f*_*eαβ*_ and *ϕ*_*eα*_ → *ϕ*_*eα*_ − *ϕ*_*eαβ*_, in order to recover Eq. (S29). We can implement analogous redefinitions if we only consider focal locus *x*_*β*_. All of our results from above then apply to each focal locus, with the appropriately defined and redefined quantities.

We can implement a similar set of definitions and redefinitions for larger numbers of focal loci. We note that, using these definitions, we have a total of 2*FE* + 2*FEn* parameters: *FE* terms 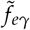, *FE* terms *f*_*eγ*_, *FEn* terms *ψ*_*eγ*_, and *FEn* terms *ϕ*_*eγ*_. This is much more than the total number of data points (*En* + *FEn*). However, we also have a corresponding set of constraints (e.g. *y*_*e*_ 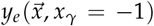 must be independent of which focal locus we consider). Using these constraints, it is straightforward to compute all of the parameters relevant for each focal locus directly from the data. We can then compute all of the relevant variances and covariances (e.g. 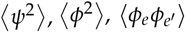, etc), and use these to compute expected patterns of global epistasis and pleiotropy for each focal locus as described above.

#### Connected Architecture and optimization problem

In this section, we describe the process used to fit parameters in the connectedness model from the background fitness and fitness effect data.

##### Optimization problem

Eq. (18) in the main text outlines the relationship between measured quantities (i.e. the variances and covariances of *ψ*^′^ *s*), and *ϕ*^′^ *s*, and the model parameters which describe interaction structure (i.e. the *w*^′^ *s* and *ν*^′^ *s*). Our goal here is to infer the latter from the former. It is useful to compile these variances and covariances, along with the isolated additive effects, into matrices that can be utilized in an optimization framework. We tabulate them here:

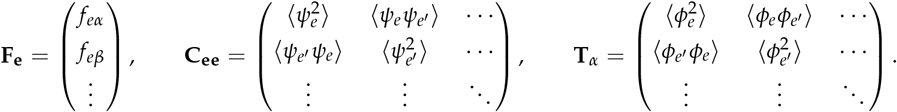

Here the **F**_**e**_ vector contains the isolated additive effects of the focal loci, **C**_**ee**_ encodes the correlation between the background loci interactions between all environments, and **T**_*α*_ represents the correlation between interaction terms of a focal locus in different environments (note there is one matrix **T**_*α*_ for each of the *F* focal loci).

##### Obtaining matrices from data

These matrices consist of *E × F* values of *f*_*eα*_, *E* values of *ψ*_*e*_, and *F × E* values *ϕ*_*eα*_, each of which is found directly from data. We start with the generalized multi-locus fitness description from Eq. (S29) in the main text,

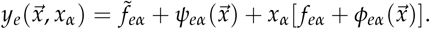

At each focal locus, we have the fitness values with and without the focal mutation,

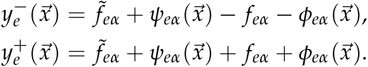

The isolated additive effects of each focal mutation are found from the means over all background genotypes, 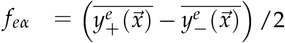. Subtracting off the means we can find the *ψ*^′^ *s* and *ϕ*^′^ *s*,

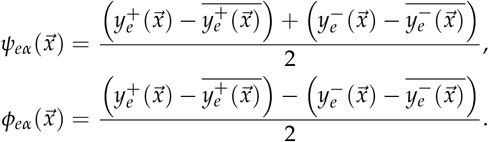

This gives us *E × F* values of 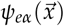 and 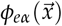. However, each background 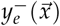 is the same for all focal loci mutations. We want to have one description of the background fitness variance for each environment. Therefore, to find *ψ*_*e*_ from each *ψ*_*eα*_ term, we change basis using

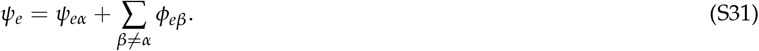

We find that the *ψ*_*e*_ values obtained from different *ψ*_*eα*_ values are in agreement. Our goal once fitting the model is to find the relationship between 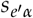 and *y*_*e*_ from the model, which is composed of components of each matrix.

##### Loss function

Given the previous definitions, we aim to minimize the following loss function:

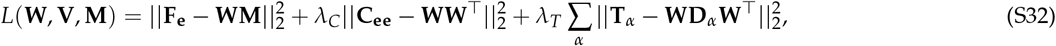

where **W** has matrix elements *w*_*eµ*_, **M** has elements *g*_*µα*_, **D**_*α*_ = diag(**V**_*α*_) and **V** contains *ν*_*µα*_ values for each locus, for all modules. Specifically, **F**_**e**_ ∈ ℝ^*E*×*F*^, **W** ∈ ℝ^*E*×*K*^, **M ∈** ℝ^*K*×*F*^, **C**_**ee**_ **∈** ℝ^*E*×*E*^, **T**_*α*_ ∈ ℝ^*E*×*E*^ for *α* = 1,.., *F* and **V** ∈ ℝ^*F*×*K*^. The hyper-parameters, *λ*_*C*_, *λ*_*T*_, and *K* (the number of modules), are fit according to the description in the following sections.

##### Optimization Process

We go through the following optimization schedule to obtain **W, V**, and **M**:

1. Least squares regression to update **M**.
2. Non-negative least squares regression to update *ν*_*µα*_’s which feed into **V**.
3. Conduct gradient steps on **W**.

Step 3 requires identifying the gradient of the loss function with respect to **W**. We can do this analytically by taking the derivative w.r.t. **W** of combinations of matrices,

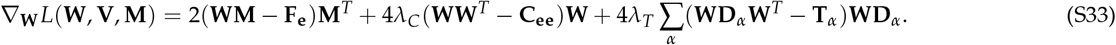

We then use this analytic gradient to take steps which change **W** with the goal of minimizing loss,

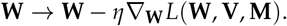

##### Fitting Hyperparameters

The loss function in Eq. (S32) establishes two sets of parameters: the model parameters **V, W**, and **M**, and hyper-parameters *λ*_*C*_, *λ*_*T*_, and the number of modules, *K*, where *K* is implicit in the dimension of matrices **W** and **M**. To determine parameters we first identify the collection of mutations which have measurements on at least 10 backgrounds in all environments. We then split the effects of each mutation into train/validation/test subsets (80%, 10%, 10% respectively) multiple times (the effects of the same mutations on different backgrounds are sorted into different subsets). For each split, we build numerical targets (F, C, T) and then fit the model using a grid of different hyperparameter combinations (*K,λ*_*C*_, *λ*_*T*_) which control the model complexity and regularization strength. The model’s mean squared error (MSE) on the validation data is used to rank configurations. After averaging validation results across repeated random splits, we select the best hyperparameters, retrain the model, evaluate it on held-out test data, and report mean ± standard-deviation MSEs for the three components (**F**_**e**_, **C**_**ee**_, **T**_*α*_). We find that the hyperparameters, *λ*_*C*_ = 0.01, *λ*_*T*_ = 10^6^, and *K* = 4 perform the best on the validation data set. Five different splits were used to determine the best performing combination.

##### Connectedness model predictions

In order to predict the slope and intercept of 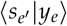 from the outputs of the connectedness model architecture, we must change basis from *ψ*_*e*_ → *ψ*_*eα*_. More specifically, due to the form of the output of the model we must find 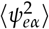 using 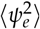. From Eq. (S31) we have

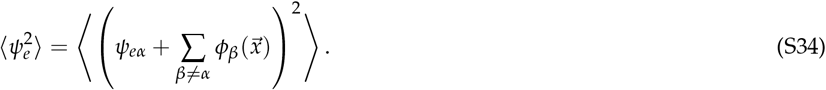

We can therefore write 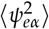 as:

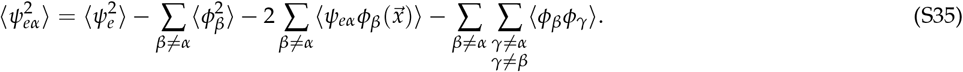

In the WRE limit, terms of the form 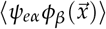 should be much smaller than the 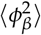 terms. The last term in the expression represents interactions which are shared by focal loci, which we also expect to be small compared to 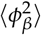. Under these assumptions we change basis using the equation 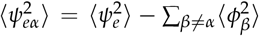. After doing so we can write the predictions for the slope and intercept in the form of Eq. (6).

#### Steady State fluctuations

We investigate three different model architectures with their own prescriptions for the relationship between loci and modules. Here we set *µ* = 1 and *µ*^′^ = 2 as well as *w*_*eµ*_ = *β*_1_, 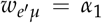, and 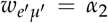 for ease of understanding in the following calculations.

In the independent case, the level of participation of a locus in different modules is drawn independently. If the modules are isolated, a given locus only participates in one module, and in the homogeneous modules case, loci will have the same level of participation in each module. In this section, using the JDFE’s previously calculated, we determine the expectations for the module 1 output when selecting in environment *e*, ⟨*σ*_*µ*_ ⟩_*e*_, as well as the three other cases: the output in module 1 when selecting in environment *e*^′^, 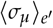, the module 2 output when selecting in environment *e*, 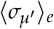, and the module 2 output when selecting in environment *e*^′^, 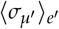.

##### Independent modules

In the case of modules in which the loci choose their degree of participation independently for each module (*µ*_*im*_ drawn independently), we want to solve the integral,

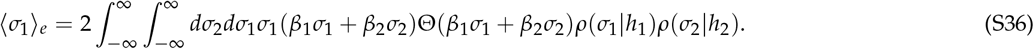

Here the heaviside function allows for only positive mutations to be selected based on the fixation probability *p*_fix_ = 2*s*_*e*_ = 2(*β*_1_*σ*_1_ + *β*_2_ *σ*_2_). We can write this as

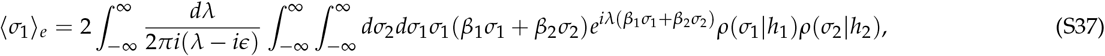

where

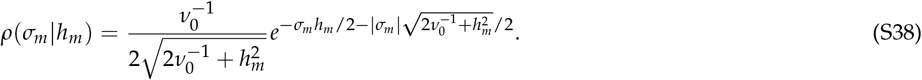

We first rescale with 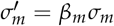

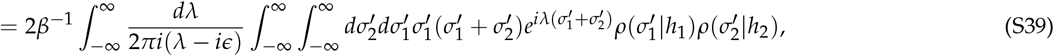

and plugging in 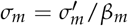 to Eq. (S38) we get

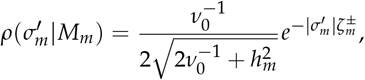

where 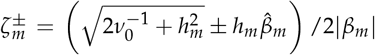. More specifically, 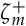 is the scaling when 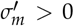 and 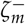 is the scaling when 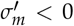 and 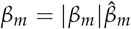. Returning to Eq. (S39) we can expand the last two integrals with

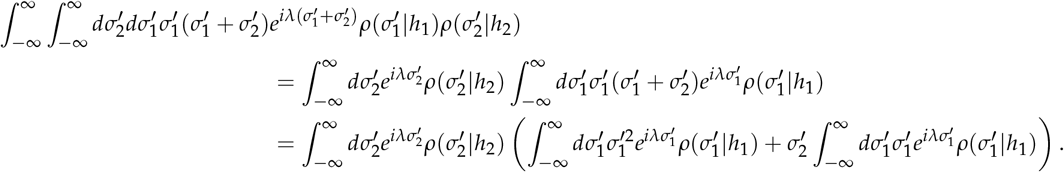

Expanding further we get

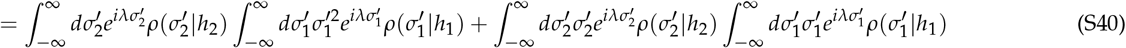

Here it is useful to move into fourier space and use the characteristic functions of the DFE’s. Using the following identities,

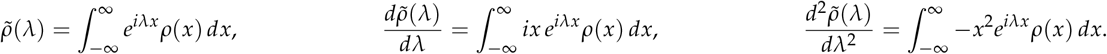

Eq. (S40) can be written as

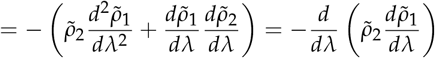

and Eq. (S39) becomes

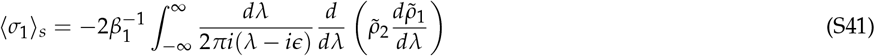

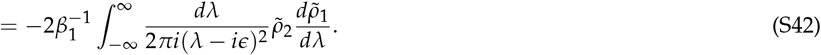

For an exponential distribution, the characteristic function is:

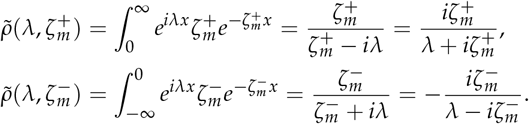

If we want 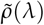 over all values of 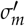 we are summing over the random variables

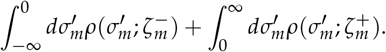

We therefore have to multiply the characteristic functions to get the general form,

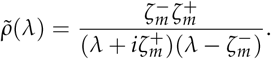

Returning to Eqn. S42 we now have

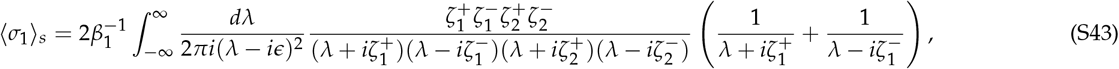

and we want to evaluate the two integrals

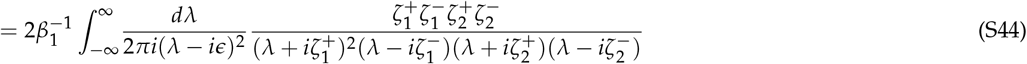

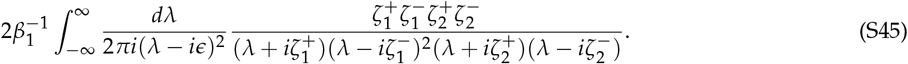

We close the contour in the lower half of the complex plane to avoid dealing with two double poles. The poles we enclose can be addressed by finding their residues. For the first integral we have:

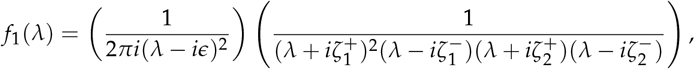

and

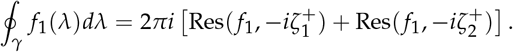

Looking at the simpler pole first:

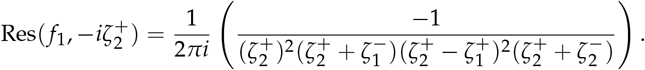

And for the repeated pole we have:

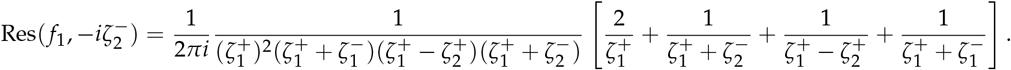

From here we can follow out similar procedures to find the result of the entire integral.

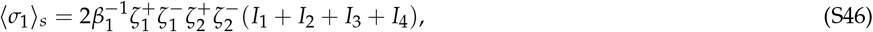

where

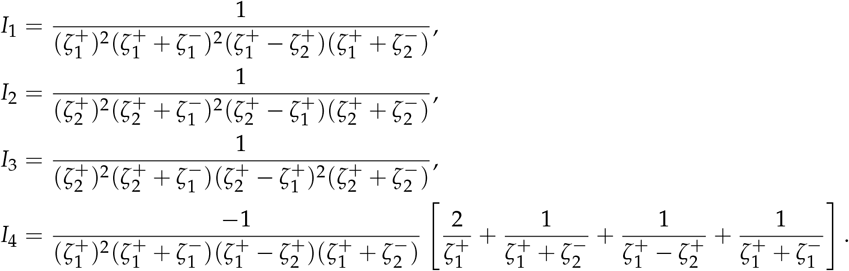

For the second module, we are instead solving the integral

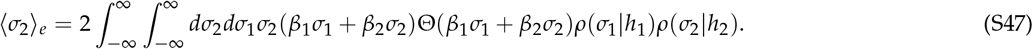

The structure of the integral remains the same. To adjust the result we just have 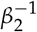 in front and we replace 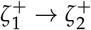 and 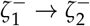. We can use these two results to find the fitness effect in the first environment ⟨*s*⟩_*e*_ = *β*_1_⟨*σ*_1_⟩_*e*_ + *β*_2_⟨*σ*_2_⟩_*e*_.

If instead we select in the second environment, we now care about *α*_1_ and *α*_2_. We must redefine our scaling variables using 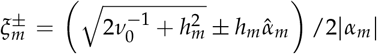 and the expectation for the modular outputs when selecting in environment *e*^′^ take the same form as environment *e* with this scaling replacement and *β* → *α*.

##### Isolated modules

When loci are restricted to participating in only one module we want to evaluate the integral

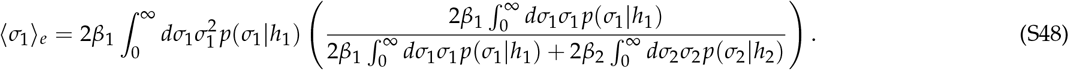

This is the expectation of the effect of the mutation given that the locus is in module 1 multiplied by the probability that a mutation will fix in module 1. This tells us that as *h*_1_ increases the rate of fixable mutations in module 1 decreases and it is more likely that module 2 will contain the mutation that fixes. Writing each type of integral out, we have:

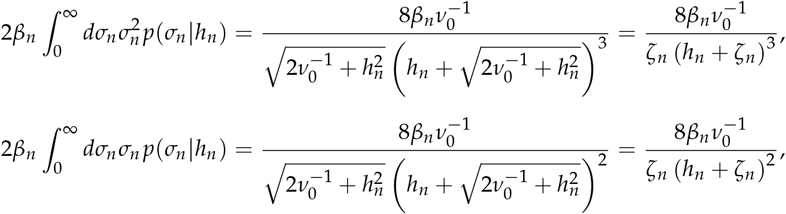

where 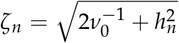. Therefore we have:

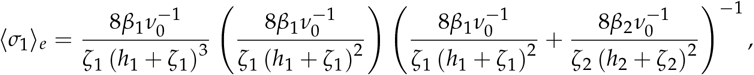

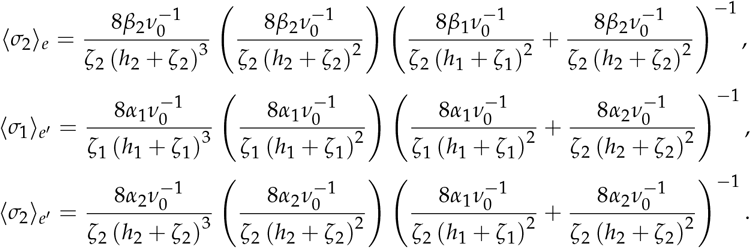

If we want to find at what values of *h*_1_ and *h*_2_ we will reach steady state, we can ask when the two modules themselves reach steady state. This will occur when

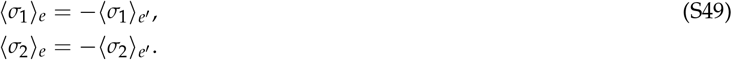

The previous calculation is the result of both environments caring positively about both modules. This creates no tension in the system and we will not observe a steady state – selection will continue to improve both environments.

If instead, an environment cares negatively about one of the modules, the fixation probability will now focus on the deleterious mutations which exist in that module – the side of the DFE which is less than zero. Since the modular DFE is generally non-symmetric (especially when a module is very weakly or strongly adapted), we can not simply apply a negative sign to the integral from 0 to ∞. Instead we we have to perform the integral from −∞ to 0.

If we consider the case where the *β* vector exists in quadrant I and the *α* vector is in quadrant III, we have the following calculation for the expectation of the effect on module 2 output when we are selecting in environment 1:

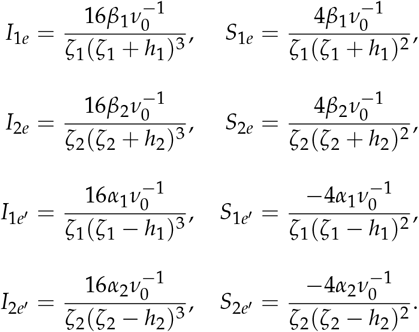

We then have the following expressions for 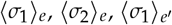, and 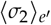,

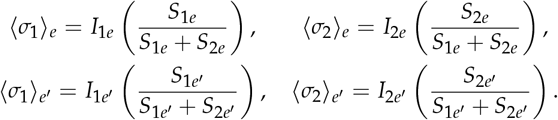

Since each locus is isolated to one module, the system can only be put under tension at the level of module dependence. Therefore, a steady state is reached in the system only when each module has opposing dependencies: one environment vector is in QII and the other is in QIV or QI and QIII, respectively.

##### Homogeneous modules

In the homogeneous modules model, the locus participation is the same for both modules, *µ*_*im*_ = *µ*_*i*_. We again write out the expectation of the output of module 1 when selecting in environment *e* as

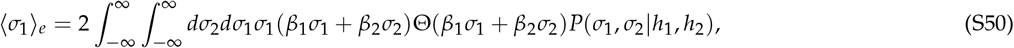

where the JDFE is defined by the integral

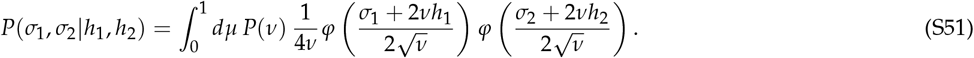

We once more assume the distribution of variance fractions is exponential, 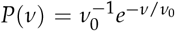. Defining the fitness effect in environment *e* as *s* = *β*_1_ *σ*_1_ + *β*_2_ *σ*_2_ and substituting in the JDFE, Eq. (S50) becomes

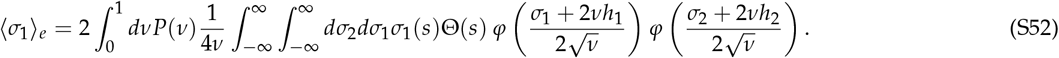

Now we can recognize the integrals over *σ*_1_ and *σ*_2_ against the two standard normals can be written as the expectation over the conditional Gaussian pair *σ*_1_, *σ*_2_ | *µ* for a fixed value of *µ*,

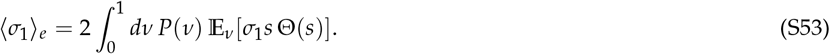

Although *σ*_1_ and *σ*_2_ are conditionally independent Gaussians given *µ*, the presence of Θ(*s*) couples them inside the integrand. So the double integral over *σ*_1_, *σ*_2_ collapses into a single joint expectation, E_*ν*_ [*σ*_1_ *s* Θ(*s*)], taken over the pair (*σ*_1_, *σ*_2_). By the law of total expectation, 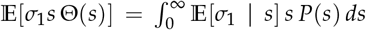 *ds*, where *P*(*s*) is the marginal density of *s* given *µ*. If we define *F* = *β*_1_ *h*_1_ + *β*_2_ *h*_2_ and 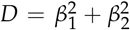, then we can write out the means and variances of *σ*_1_ and *s* as E[*σ*_1_] = −2*νh*_1_, E[*s*] = −2*νF*, Var(*s*) = 4*νD*, and Cov(*σ*_1_, *s*) = 4*νβ*_1_. Using these, the conditional is

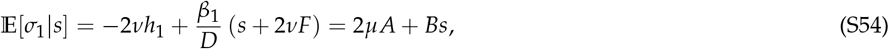

where *A* = *Fβ*_1_/*D* − *h*_1_ and *B* = *β*_1_/*D*. With this expression, the expectation in Eq. (S53) can be written as

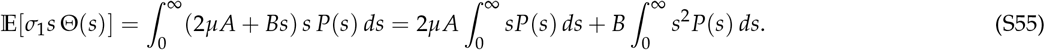

Since *s*|*µ ~ N* (−2*νF*, 4*νD*), the truncated moments are

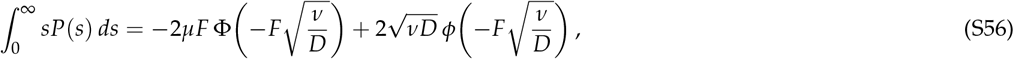

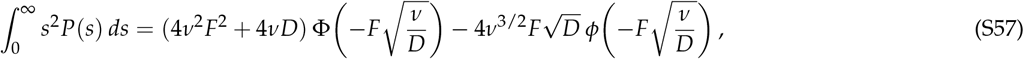

where φ(*x*) and *ϕ*(*x*) denote the standard normal cumulative distribution function and probability density function, 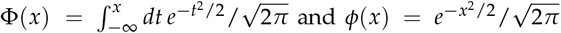, respectively. Substituting these truncated moments into Eq. (S55), and then into Eq. (S53), leaves only integrals over *ν*. All resulting terms can be expressed as linear combinations of three basic integrals. Defining *r* = 1/*µ*_0_ and *q* = *r* + *F*^2^/2*D*, these integrals are

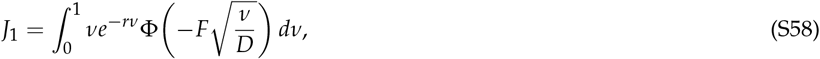

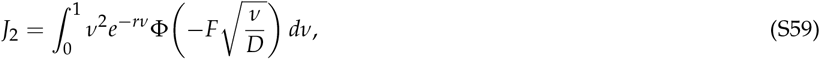

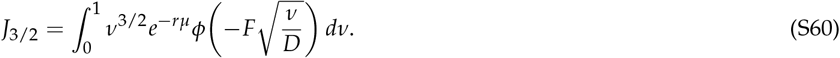

And they evaluate to

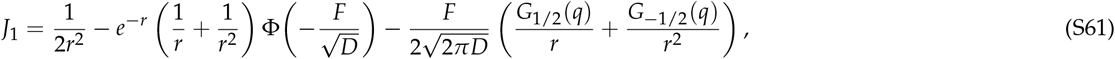

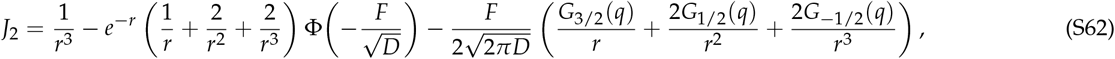

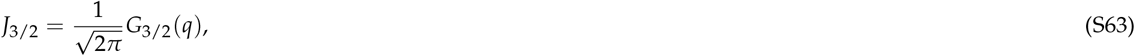

where 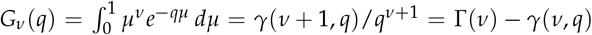 is the upper incomplete gamma function. Finally we can express the expectation of the modular fitness effect when selecting in environment *e* in terms of these solutions and previously defined parameters as

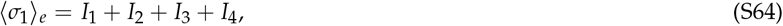

where

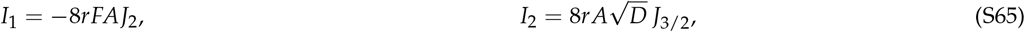

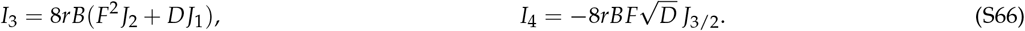

For the other three modular expectations the solutions come in similar forms with the defined variables. If instead we are finding the expectation of *σ*_2_, *A* → *A*^′^ = *Fβ*_2_/*D* − *M*_2_ and *B* → *B*^′^ = *β*_2_/*D*. And when selecting in environment *e*^′^ the *β*^′^ *s* are replaced by *α*^′^ *s*.

#### Experimental Materials and Methods

##### Growth assay details

Frozen stocks were revived by completely thawing all segregant libraries, mixing well, and transferring 8 µl of frozen stock into 120 µl of YPD+Hyg+Nat (300 µg /ml HygB (Gold Biotechnology Inc #H27010) and 20 µg /ml cloNAT (Gold Biotechnology N-500-100)) in a 96-well flat-bottomed plate. Each segregant was kept separate in a well, and is identified by the well number. These were allowed to grow for 36 hours, shaken at 30°C, until saturation. After this initial grow-up, we added 2 µl of saturated culture to 126 µl of chosen media (YPD, YNB, or Glyc) in flat-bottomed, unshaken 96-well plates, and allowed to saturate in the appropriate temperature. After 24 hours of growth, we marked this as time point zero, sampled for sequencing, and conducted the next transfer. Each segregant occupies the same well in all plates, for all transfers. This 1 : 2^6^ transfer was conducted every 24 hours for 8 days, leading to 42 generations of growth for the assay.

**Table S1:** Growth Conditions and Media Details. All media also contained 1 µg/ml ampicillin. Reagents used: Yeast Extract (VWR #76628-828), Bacto Peptone (VWR #76628-742), Dextrose (Fisher Sci #BD-215530), Yeast Nitrogen Base (VWR #90004-400), Glycerol (VWR #BDH1172), Ampicillin (Sigma-Aldrich #A9518).

| Growth Condition | Details of Media |
| --- | --- |
| YPD at 23°C | 1% yeast extract, 2% Bacto peptone, grown on bench |
| YPD at 30°C | 1% yeast extract, 2% Bacto peptone, 2% dextrose, grown at 30°C and 40% humidity |
| YPD at 37°C | 1% yeast extract, 2% Bacto peptone, 2% dextrose, grown at 37°C and 50% humidity |
| Glycerol at 30°C | 1% yeast extract, 2% Bacto peptone, 0.5% dextrose, 3% glycerol, grown at 30°C and 40% humidity |
| YNB at 30°C | Yeast Nitrogen Base w/o amino acids, 2% dextrose, grown at 30°C and 40% humidity |

For pelleting and sequencing, we took 45 µl of each segregant and mixed them into a single pellet at each transfer. We could do this since each barcode corresponds to a specific plasmid library (which identifies the segregant) and edge (which identifies the mutation). This allows us to pool before DNA extractions and sequencing, and demultiplex data during analysis. Each pellet was spun down, the supernatant discarded, and the pellet frozen at −20°C until it was prepared for sequencing. Three different replicate assays were conducted for 5 different growth conditions, leading to a total of 15 plates being maintained for each day of the transfer. All 15 replicates were assayed at the same time, though each replicate used a different batch of growth media.

##### DNA Extractions for barcode sequencing

Each mixed pellet was approximately 4 mL. A full cell pellet was incubated at 37°C in 100 µl zymolyase lysis buffer (5 mg/mL Zymolyase 20T (Nacalai Tesque), 1 M sorbitol (Sigma-Aldrich S1876), 100 mM sodium phosphate pH 7.4, 10 mM EDTA (Thermofisher B49), 0.5% 3-(N,N-Dimethylmyristylammonio)-propanesulfonate (Sigma-Aldrich #40772), 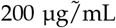 RNAse A (QIAgen #19101), and 20 mM DTT(Goldbio) for 45 minutes. 400 µl of a binding buffer with guanidine thiocyanate (4 volumes of 100 mM MES pH 5 (Goldbio #M-090), 4.125 M guanidine thiocyanate(Goldbio #G-210), 25% isopropanol, and 10 mM EDTA was added, well mixed, and spun down to remove any cell debris. The supernatant was added to a silica column (IBI Scientific #IBI-IB47207) to allow the DNA to bind to the column, and the flowthrough was discarded. After binding, the columns were washed with 400 µl of a first wash buffer (10% guanidine thiocyanate, 25% isopropanol, 10 mM EDTA) and then 600 µl of a second wash buffer (80% ethanol, 10 mM Tris pH 8 (Corning #46-031-CM)). An extra spin down step for 3 minutes allowed any remaining ethanol to evaporate, for better downstream processing. All samples were eluted into 50 µl of molecular-grade water, yielding between 20–30 ng/µl of gDNA, depending on growth conditions.

##### Sequencing Library Preparation and PCR protocols

For each sample, we performed round 1 reactions which enable each amplicon to be tagged with a UMI. Each forward primer also has a random 7 base-pair region. Each amplicon should have a unique UMI; repeats may occur due to PCR bias and are removed during data analysis. There are also 12 different forward primers to generate an offset in each amplicon, creating the diversity needed for a patterned sequencing flow cell.

To set up the reaction, we combined 15 µl gDNA, 25 µl 2X Kapa HotStart HiFi MM, 3 µl 10 µM TnRS1 primer, 3 µl 10 µM TnFX primer, and 4 µl molecular-grade water. We ran the PCR protocol: 1) 95°C for 3:00, 2) 98°C for 0:20, 3) 60°C for 0:30, 4) 72°C for 0:30, GO TO step 2 for 4 cycles, 5) 72°C for 1:00. This was purified with a left-sided cleanup using Aline magnetic beads (0.85x ratio) and eluted in 30 µl of molecular-grade water. All primers can be found in Johnson et al. [2019] and were ordered from Integrated DNA Technologies (IDT).

To add Illumina adapters, this cleaned-up PCR product was amplified and dual-tagged in round 2 using Nextera primers. The round 2 reaction combined 25 µl purified PCR 1 product, 8 µl molecular-grade water, 10 µl Kapa HiFi Buffer, 1 µl KAPA HiFi HotStart DNA Polymerase, 1 µl dNTP mix, 2.5 µl 10 µM N7XX primer (Nextera), and 2.5 µl 10 µM S5XX primer (Nextera), and used the PCR protocol: 1) 95°C for 3:00, 2) 98°C for 0:20, 3) 61°C for 0:30, 4) 72°C for 0:30, GO TO step 2 for 25 cycles, 5) 72°C for 2:00. After visualization on an agarose gel, we cleaned the prepped libraries using another left-sided bead cleanup and Aline magnetic beads (0.85x ratio), eluting in 50*~ µ*l of molecular-grade water.

All libraries were quantified using a fluorescent plate-based assay on the SpectraMax i3 and the AccuGreen ™ High Sensitivity dsDNA Quantitation Kit (Biotium #31066). All prepped libraries were pooled by weight. A final 0.8x-0.2x left and right sided cleanup using magnetic Aline beads was performed on the pool to remove any remaining small and large fragments, and fragment size distribution was checked on an Agilent Tapestation 4200. The sequencing was done on 2 separate runs on NovaSeq X Plus 10B lanes (Illumina) using paired end, 2×150 reads.

##### Inferring mutational fitness from sequencing data

The mutational fitness values were inferred using the process outlined in Johnson et al. [2019]. We measure the log-frequency slope between successive time-points (each time-point must have a minimum of 4000 total counts). Since the neutral barcodes are already known, we can use the median log-frequency slope of the neutral set of barcodes to make the mean fitness correction to these slopes. After excluding assays with less than 5 neutral barcodes (with at least 30 counts each) or less than 3 usable time points, we find fitness effects by scaling values, and averaging over all time intervals that pass our thresholds in the assay.

Since each mutation or “edge” is redundantly barcoded, we can exclude outliers in a group of barcodes for a particular mutation by using a log-likelihood test of belonging to the group. For this process, each group must contain at-least three redundant barcodes. We measure the fitness effect for all barcodes of a mutation, and combine counts from all barcodes that are within a 0.01 of the median fitness effect for that mutation (called the ‘reference’ for that pool. For a barcode where the fitness effect is further than 0.01 from this median, we use the combined counts and the counts of that particular barcode and find the log-likelihood of 1) barcode and combined ‘reference’ have the same fitness, and 2) the barcode and combined ‘reference’ have different fitnesses, modeling non-stochastic changes in frequency of lineages, where the log-likelihood of the data at each time-point *t*, is given by a frequency prediction *f*_*k,t*_ and multinomial number of reads *n*_*k,t*_ for each of *k* lineages. We exclude the barcodes with a LL ratio above a heuristic cutoff, as determined by Johnson et al. [2019].

After exclusion of these outliers, we can further reduce the noise in measurements by collapsing redundant barcodes per mutation into combined barcodes (cBCs). These cBC’s have lower per-barcode noise due to having higher counts, but still preserve replicate fitness measurements. We repeat the fitness measurement protocol on these cBCs, and then average to get the fitness effect of a mutation against a particular genetic background. We set a threshold of a minimum number of three cBC’s for a mutation to be included. Johnson et al. [2019] have also excluded a few strains and mutations by visual inspection, those are excluded from our analysis too.

Once all three replicate fitness effect values are found we only include mutations which agree among these replicates. We first exclude any mutations which only have measurements in one replicate. For each mutation with two or more available replicate estimates, the maximum pairwise disagreement, Δ_max_ = max (*s*_rep1_, *s*_rep2_, *s*_rep3_) − min (*s*_rep1_, *s*_rep2_, *s*_rep3_), was computed. Mutations with Δ_max_ ≤ 0.05 were considered reproducible and retained, using the average of all available replicates. For mutations with three replicates whose overall disagreement exceeded 0.05, we tested whether any two of the three replicates agreed with each other within 0.05; if so, the mutation was retained using the mean of the two agreeing replicates, with the disagreeing replicate excluded. Mutations for which no pair of replicates agreed within 0.05, or for which only two replicates were available and disagreed beyond 0.05, were excluded from downstream analysis. The 0.05 threshold was chosen based on the empirical distribution of replicate disagreement across mutations (median Δ_max_ of 0.01–0.02, with the great majority of mutations disagreeing by less than 0.05), such that the filter removed a minority of poorly-reproducing measurements while retaining the bulk of the dataset.

##### Background Fitness Estimation

To set up the flow cytometry assays, we revived frozen stocks of the strain backgrounds without any mutations. 2*µ*l of fully thawed culture was added to 126 *µ*l YPD and grown till saturation in shaken, flat bottomed plates. In addition, the florescent reference strain containing a mCitrine marker (LK3-B08 from Jerison et al. [2017] was also revived and grown to saturation in flat bottomed plates. After an initial growup, the reference strain (LKB08) and the parental strains were mixed equiproportionaly, and 2 *µ*l of this was transfered to fresh YPD media. A 6 generation assay (to match the assays for the fitness effects) was conducted by transferring 2 uL of the co-culture to fresh media. The saturated co-culture was the initial time point, and the end point was taken after a further 6 generations. At each time point, 10 uL of saturated culture was transferred to 116 uL of cold, sterile PBS, and kept at 4C before being put through the flow cytometer.

Samples were run on the 4 laser LSR Fortessa and a 5 laser FAC A3 Symphony. All Glycerol and YNB samples (except rows E,F,G,H from YNB Rep1 at timepoint 1) were run on the Symphony, and all other samples (including rows E,F,G,H from YNB Rep1 at timepoint 1) were run on the Fortessa. For each well, 20,000 cells were counted, and the cytometer had a wash step between each well.

For the analysis of flow cytometry data, the first and last 10% of counted events were discarded. Potential doublet events (where the FSC-H to FSC-W ratio was above a threshold of 1.2) were also discarded. The exact ratio used made little difference to our calculations. A Gaussian mixture model was fitted to the data to separate cells into two types (fluorescent reference vs wild type parental strain) and each type was counted separately. Certain counts were manually fixed by eye if the GMM fit was deemed incorrect. Once counted, fitness against the reference florescent strain was measured by calculating 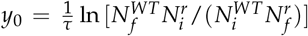 where *τ* = 6 and values in the natural log are final and initial counts of the reference and WT strains. For YNB, a 12 generation assay was conducted, and hence *τ*_*YNB*_ = 12.

**Figure S10:**
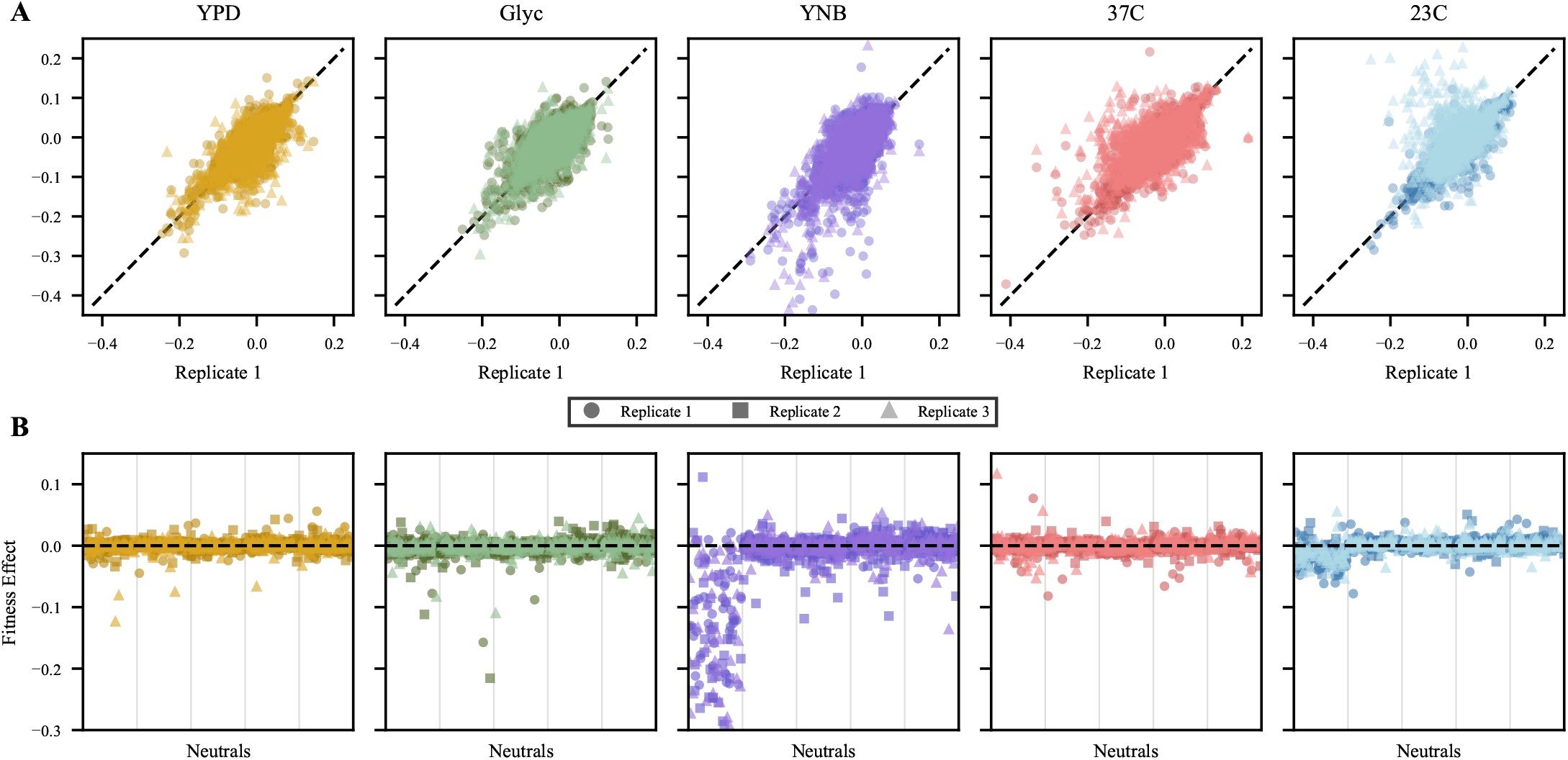
Replicate – replicate comparisons and neutral analysis of mutational fitness effects. **(A)** Mutational fitness measured in Replicate 1 vs. Replicate 2 (darker squares) and 3 (lighter triangles). **(B)** Mutational fitness measurements for the five marked neutrals in all three replicates. Each replicate can be differentiated by the color and shape of the markers. The first neutral in YNB exhibited a large spread of fitness effects and for this reason was discarded as a neutral.

For each background and environment, the difference between replicate fitness estimates was computed, Δ_e_ = *y*_*e*,rep1 −_ *y*_*e*,rep2_, and the standard deviation of this difference, *σ*_Δ_, was calculated across all backgrounds within that environment. Background-replicate pairs whose absolute difference exceeded two standard deviations of the pooled replicate-difference distribution, |Δ_e_| > 2*σ*_Δ_, were considered technical outliers and excluded from further analysis. For all retained background-replicate pairs, the final fitness estimate was recalculated as the mean of the two replicates following outlier removal. We also limit all environments to background fitness values between −0.3 and 0.3 which removes at most three data points per environment and the 37° *C* environment to points above −0.08 due to mutations made in KRE33, a gene involved in small ribosomal subunit assembly, having outsized effects important to this environment [Jerison et al., 2017]. The background fitness values have good inter-replicate correlations and can be found in Fig. S11.

**Figure S11:**
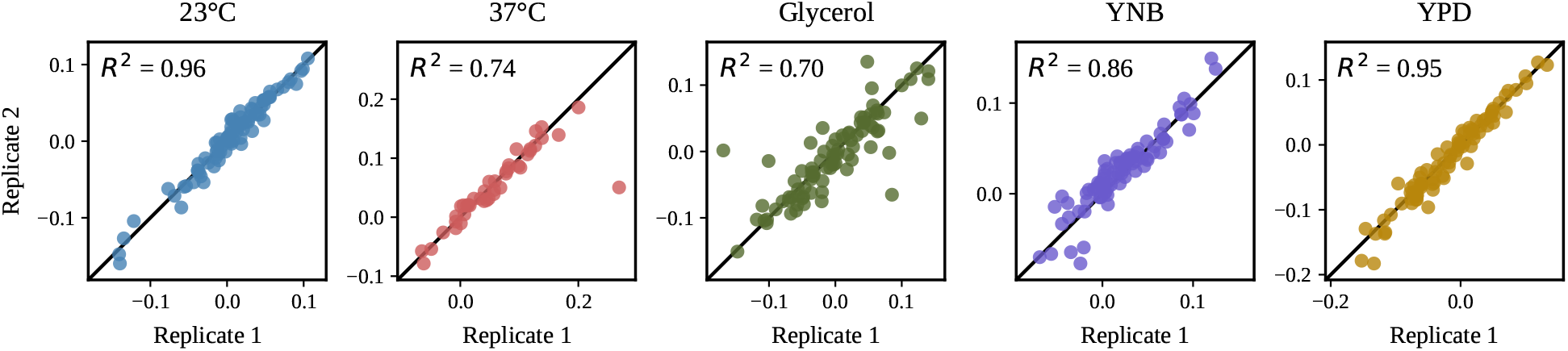
Background fitness replicate – replicate comparisons. Inter-replicate correlations of background fitness measurements through flow cytometry after outlier exclusion for all five environments. Replicate-replicate environmental correlations before outlier exclusion are included in data parsing scripts (see Code and Data Availability).

## Notes

### Competing Interest Statement

The authors have declared no competing interest.

### Summary of Updates

Corrected LaTeX formatting in the bioRxiv abstract. The manuscript content is unchanged.

